# Stress-Survival Pathway Profiling Reveals MCM10 as a Candidate Biomarker of Hispanic Colorectal Cancer Disparities

**DOI:** 10.64898/2026.01.06.698057

**Authors:** Md Zahirul Islam Khan, Urbashi Basnet, Soumya Nair, Aditi Kulkarni, Frances A. Rangel, Angel Torres, Anamika Basu, Igor C. Almeida, Taslim Al-Hilal, Brian I Grajeda, Sourav Roy

**Affiliations:** Department of Biological Sciences, University of Texas at El Paso, El Paso, TX 79968, USA; The Border Biomedical Research Center, University of Texas at El Paso, El Paso, TX 79968, USA; Science and Math Division, Copper Mountain College, Joshua Tree, CA 92252, USA; Departments of Molecular Pharmaceutics and Biomedical Engineering, University of Utah, Salt Lake City, UT 84112, USA

**Keywords:** Colorectal cancer, stress survival pathway, Hispanics, Non-Hispanic whites, health disparities, biomarkers

## Abstract

**Background:** Colorectal cancer (CRC) remains the second leading cause of cancer mortality in the U.S., with significant racial and ethnic disparities in incidence, survival, and mortality rates. The rising incidence of early-onset CRC and the frequent presentation at advanced stages of CRC among the Hispanics makes this an important ethnic group to study. A deeper understanding of the complex interplay between molecular, genetic, and environmental factors is critical for developing targeted therapies to reduce the prevalence in specific racial and ethnic groups, such as Hispanics. This study explores the role of stress-survival pathway (SSP) genes in early-onset and late-stage CRC, primarily focusing on disparities between Hispanics and Non-Hispanic Whites (NHWs). Additionally, this study investigates the role of MCM10, in contributing to CRC disparities among the Hispanic populations with an emphasis of early-onset and late-stage CRC Hispanics.

**Methods:** One of our previous studies had identified some SSP protein coding genes associated with CRC. The transcript and protein level expressions of these genes were validated in CRC cell lines, cDNA arrays, and tissue microarrays, using qRT-PCR and immunohistochemistry, respectively. The transcript level expressions of differentially expressed SSP genes were further evaluated in Hispanic and NHWs tumor tissues, including early onset and late-stage cohorts. Additionally, we performed cellular, physiological, and functional assays to explore the role of MCM10 in tumor progression, before and after siRNA mediated knockdown of MCM10. High-throughput transcriptomic and proteomic analyses were performed to reveal the underlying molecular mechanism.

**Results:** We observed the differential expressions of twelve SSP genes in CRC cell lines, at the transcript level; that of ten genes in early and late stages using cDNA arrays, and nine genes at the protein levels using tissue microarrays and immunohistochemistry. Hispanic CRC samples showed differential expression of all nine SSP genes compared to NHWs, in early-onset, and late-stage CRC, with all genes being upregulated other than CDK4 and PRDX4 in late-stage CRC and CDK4 in early-onset CRC. Our functional study demonstrated that MCM10 knockdown in CRC cell lines significantly reduced cell proliferation, growth, invasion, and migration by inducing cell cycle arrest, apoptosis, and reactive oxygen species pathways. The integrative RNA-sequencing and proteomics study identified that PPFIA1 could be associated with MCM10 mediated cancer progression. The role of MCM10 and PPFIA1 in cancer progression has also been validated by CRISPR-Cas9 mediated MCM10 knock out, using a Hispanic CRC patient derived organoid.

**Conclusion:** The differential expression of SSP genes suggest a potential molecular contribution to CRC disparities in the Hispanic population. The essential oncogenic role of MCM10 and its axis with PPFIA1 was identified; this could lead to a new avenue for therapeutics targeting the MCM10-PPFIA1 axis, thereby, combating early-onset and advanced-stage CRC in this high-risk ethnic group.

*Graphical Abstract:* 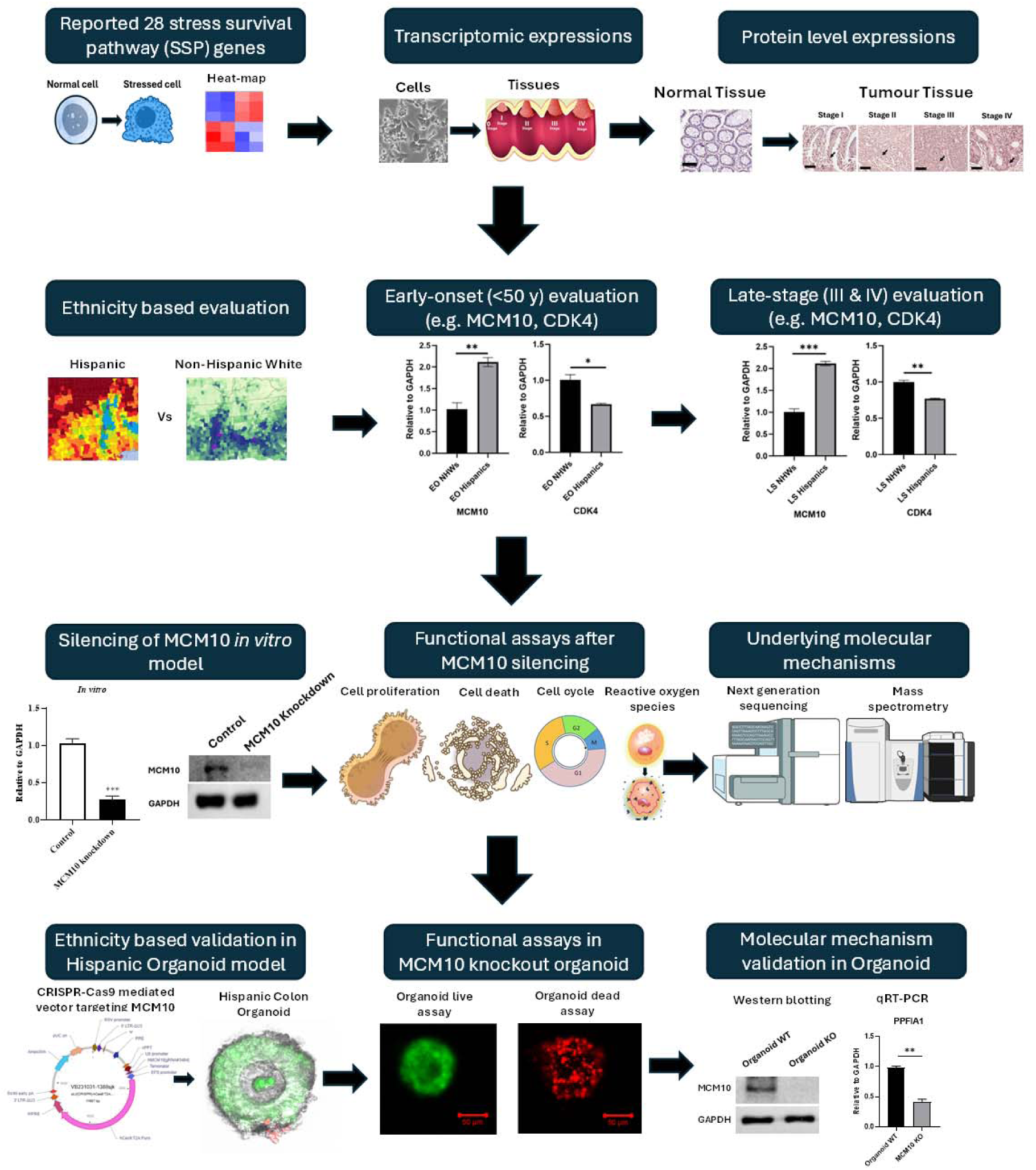

## 1. INTRODUCTION

Colorectal cancer (CRC) is the second most-deadly cancer, even though it stands at number four in terms of cancer cases by sites^1^. It is also one of the most preventable and treatable cancers when detected at an early stage^2^. The global burden of the disease is anticipated to rise substantially, with an estimated 3.2 million new cases and 1.6 million deaths annually by 2040, representing increases of 63% and 73%, respectively^3^. China, the United States (U.S.), and Japan are the countries most heavily impacted by CRC, with Russia and India also facing substantial disease burden^4^. In the U.S. population CRC incidence and mortality rates have declined steadily, primarily due to advancements in screening methods, early detection, and treatment options^5^. However, despite these improvements, CRC incidence, survival, and mortality rates vary significantly across different racial and ethnic groups^6^. The decline has been much less significant among Hispanics compared to non-Hispanic Whites (NHWs)^7^.

There has also been an alarming rise in the incidence of early-onset CRC in the U.S., with current trends projecting that these cases could be doubled by 2030^8^. This trend is particularly worrisome, as early-onset CRC is often associated with higher pathological grade and an increased risk of recurrence and metastasis^9,10^. In addition to these characteristics, early-onset CRC tumors are molecularly and pathologically heterogeneous in patients from different geographical locations and specific ethnic groups^11^. Incidences of early-onset CRC (defined as diagnosis before age of 50) among Hispanics (16.5%) are significantly higer compared to NHWs (8.7%)^12^, underscoring a unique and pressing inequity in CRC burden within the Hispanic population. Several studies have hypothesized that unhealthy diet, family history, hyperlipidemia, male sex, alcohol consumption, tobacco use, sedentary lifestyle, obesity, and diabetes may contribute to the of early-onset CRC^13,14^. This disparity is further compounded by the fact that Hispanics contribute to a higher proportion of later-stage diagnoses compared to NHWs (32% vs. 19% diagnosed at Stage IV), which likely accounts for the inferior age-adjusted survival in Hispanics, following CRC diagnosis^15^.

Addressing disparities in CRC among Hispanics, especially the rising incidence of early-onset CRC and the frequent presentation at advanced stages, is essential to improving outcomes in this underserved population^16,17^. A deeper understanding of the complex interplay between molecular, genetic, and environmental factors is critical for developing targeted therapies to reduce the prevalence and implementing ethnicity-specific screening and treatment strategies, particularly in specific racial and ethnic groups, such as Hispanics, where CRC represents a growing public health challenge^18^.

In normal colonic epithelium, cells mount a coordinated response to stress that sustains a dynamic equilibrium between apoptosis and cell proliferation, which is essential for maintaining cellular homeostasis. However, disruption on this balance contributes to the initiation and progression of CRC^19^. Similarly, chronic stress is recognized as a significant risk factor for CRC progression, as activation of stress-survival pathways (SSP) can promote carcinogenesis and therapeutic resistance^20^. Reactive oxygen species (ROS), which are metabolic byproducts of cellular processes, function as redox signaling messengers that regulate numerous aspects of cellular activity and contribute to the maintenance of homeostasis^21^. Oxidative stress arises when there is an imbalance between ROS production and the antioxidant defense system. This imbalance contributes not only to CRC development but also to impaired responses to therapeutic interventions^22,23^. The progression from normal colorectal epithelium to adenoma and ultimately to carcinoma involves a cascade of genetic and epigenetic alterations, including inactivation of key tumor suppressor genes, activation of proto-oncogenes, disruption of the extracellular matrix, DNA mutations, and aberrant DNA methylation patterns^24^. It is believed that oxidative stress and apoptosis are closely interconnected physiological processes that play a critical role in CRC initiation and progression^25,26^. In a previous study from our laboratory, we identified a set of SSP-related protein-coding genes associated with CRC through in-silico analysis of multiple cancer databases^27^.

In this study, we present novel findings highlighting the role of SSP genes in contributing to disparities in early-onset and late-stage CRC among Hispanics. Given the rising incidence of early-onset CRC in this population and the frequent occurrence of late-stage diagnoses, we analyzed the expression of SSP genes in Hispanic and NHW patients. We further investigated the functional role of one SSP gene, *minichromosome maintenance 10* (MCM10), in CRC progression. Knockdown of *MCM10* suppressed CRC carcinogenesis by reducing cell proliferation, invasion, and migration, and by modulating cell cycle, apoptosis, and ROS pathways. We also examined the contribution of *MCM10* to early-onset and late-stage CRC disparities in Hispanics. and validated its functional relevance using CRC patient-derived organoids. Overall, this study offers novel insights into the contribution of SSP genes to CRC progression and racial and ethnic disparities, with particular emphasis on the role of *MCM10* in Hispanic CRC.

## 2. MATERIALS AND METHODS

### 2.1. Patient samples collection and RNA extractions

This study utilized CRC tissue along with their normal-adjacent tissue (NAT) samples from both U.S. and non-U.S. sources, with particular focus on individuals from Hispanic and NHW. The tissue specimens were procured from certified commercial biorepositories, including Reprocell Inc. (MD, USA), Proteogenex Inc. (CA, USA), Tissue for Research (Newmarket, UK), BioIVT LLC (TX, USA), and iSpecimen (MA, USA). Among these, Proteogenex Inc. provided high-quality, pre-isolated total RNA. For all remaining vendors, fresh-frozen tissue samples were acquired, and total RNA was extracted in-house using the Quick-RNA Miniprep Plus Kit (Zymo Research Corporation, CA, USA), following the manufacturer’s protocol. This approach ensured consistency and quality control across diverse sample origins.

### 2.2. Cell lines and transfection

Seven cell lines were obtained from American Type Culture Collection (ATCC, Virginia, USA), including a normal colon cell line (CCD841 CoN) and six CRC cell lines (SW1116, SW480, SW620, SW48, Caco-2, and WiDr). All the cell liens cultured and maintained according to the ATCC guidelines. To achieve MCM10 knockdown, three small interfering RNA (siRNA) targeting MCM10, and negative controls (scrambled) were designed and purchased from MilliporeSigma (MA, USA). The transfection was performed using a lipid-based transfection reagent Lipofectamine RNAiMAX (Invitrogen Corporation, CA, USA). Among the three tested siRNAs, siRNA #1 and siRNA #2 displayed maximum MCM10 knockdown efficiency where siRNA #1 was used for further experiments. The transfection efficiency was measured using both qRT-PCR and western blot respectively.

### 2.3. Complementary DNA (cDNA) synthesis and qRT-q

The cDNA was synthesized from 1,000 ng of total RNA using the RT Master Mix (Bio-Rad Laboratories, CA, USA), following the manufacturer’s protocol. Quantitative real-time PCR (qRT-PCR) was conducted using the SsoAdvanced Universal SYBR Green Supermix (Bio-Rad Laboratories) on the CFX Opus Real-Time PCR System (Bio-Rad Laboratories). Relative gene expression levels were determined using the 2^(-ΔΔCt) method, with GAPDH serving as the in-house control.

### 2.4. cDNA arrays and tissue microarray slides (TMAs)

A commercially available Colon Cancer cDNA Array IV was obtained from OriGene Technologies Inc., (MD, USA), which included cDNA samples from 8-normal, 5-Stage I, 1-Stage II, 8-Stage IIA, 1-Stage IIIA, 6-Stage IIIB, 3-Stage IIIC, 6-Stage III, 10-Stage IV.

The protein level expression of target genes was detected using commercially available human Colon Cancer Tissue Microarrays (CRC TMAs) from US Biomax, Inc. (MD, USA) containing 5 Stage I, 39 Stage IIA, 28 Stage IIB, 24 Stage IIIB, 8 Stage IIIC, 3 Stage IV and 12 normal colon tissue, single core per case.

### 2.5. Cell proliferation, colony formation, invasion and migration assay

Twenty-four hours post transfection, both Caco-2 and SW480 cells were trypsinized, counted and seeded (10 x10^3^) in a 96-well plate. Following the CCK-8 protocol, 10□µl of CCK8 solution (MCE) was added in each well and viability was assessed by measuring formazan absorbance at 450nm in Microplate Absorbance Reader (Biotek Laboratories, WA, USA) at 0, 24, 48, and 72□hours.

Similarly, 500 cells were seeded in 6-well plate for colony formation assay to assess the cell proliferation and growth capacity *in vitro*. After the colony formation, they were fixed and stained with 0.5% crystal violet to visualize the colonies and captured & counted them under EVOS XL Core microscope (Thermo Fisher Scientific, MA, USA).

To evaluate the invasive and migratory characteristics of Caco-2 and SW480 after knocking down MCM10, we performed Transwell invasion and migration assay on 24 well cell culture inserts with an 8 μm pore size (Corning Inc., NY, USA). For the invasion assay, 1 x 10^5^ cells were seeded into the Matrigel (Corning Inc.) coated upper chamber, while for migration assay, an uncoated chamber was used. The lower chambers contained 20% FBS which played a role as chemoattractant. Cells that invaded or migrated through the membrane were fixed, permeabilized, and stained with 0.1% crystal violet. The stained cells were then further visualized and quantified using microscope EVOS XL Core (Thermo Fisher Scientific, MA, USA).

### 2.6. Cell cycle, apoptosis and ROS assay

The transfected Caco-2 and SW480 cells and their respective controls were trypsinized and fixed with 70% ethanol overnight at 4°C. Next day, RNaseA (20□μg/mL) and propidium iodine (MCE) were added to the cells and incubated for 30 min in a dark place according to the manufacturer’s protocol. Data acquisitions and processing were performed on flow cytometer (Beckman Coulter, Inc., CA, USA).

Likewise, the apoptosis was detected using Annexin V Apoptosis Detection Kit with 7-AAD (Tonbo Biosciences, CA, USA). Based on the manufacturer’s protocol, cells were treated with Annexin V and 7-amino-actinomycin (7-AAD) in Annexin V Binding Buffer (TONBO Biosciences). The apoptotic cells were then measured using Beckman Coulter flow cytometer.

The ROS levels in both CaCo-2 and SW480 cells were assessed using a fluorescence microscope with the ROS Assay Kit (ABP Biosciences, MD, USA). Briefly, cells were seeded in 48 or 96-well plates, incubated for 24 hours, and subsequently transfected with siRNA of MCM10. Furthermore, 10mM H_2_DCFDA solution was added into the cells, followed by 30 minutes incubation and immediate visualization of ROS under LSM 700 Confocal Microscopes.

### 2.7. Immunohistochemical staining and scoring

The Formalin Fixed Paraffin Embedded (FFPE) TMAs were deparaffinized in xylene and rehydrated through a serial dilution of ethanol followed by PBS. Antigen retrieval was performed using Citra-Plus solution (Biogenex Laboratories, CA, USA) in a microwave. Endogenous peroxidase activity was blocked with 3% hydrogen peroxide in 10% methanol, and non-specific binding was minimized with Universal Blocking Reagent (Biogenex Laboratories). The TMAs were incubated overnight at 4°C with primary antibodies against BCL2L1, CCNB1, CDK1, CDK4, Chk1, CSE1L, MCM10, PDCD2L, PRDX4, and FOXM1 at an optimized dilution. Next day, secondary antibodies from Santa Cruz Biotechnology, (Texas, USA) and Abcam Limited (Cambridge, England) were applied, followed by Streptavidin-coupled peroxidase. The source for primary and secondary antibodies used in this study were mouse and rabbits, respectively. The AEC solution was used as chromogen, and slides were counterstained with haematoxylin, mounted, and dried overnight. The imaging was performed with an Aperio CS2 slide scanner (Leica Biosystems, IL, USA) at 200x magnification. Staining intensity was blindly evaluated by three independent observers in a score ranging from 0 to 3 which represents negative, weak, moderate, and high protein expression.

### 2.8. Western blotting

Cells were collected with or without transfection and lysed in an appropriate amount of RIPA lysis buffer from MedChemExpress LLC. (MCE, NJ, USA). Furthermore, proteins were quantified using BCA Protein Assay Kit (Thermo Fisher Scientific, MA, USA) and performed western blot following previously described optimized protocol^28^. Briefly, 30-50µg of proteins from each sample were separated into 8% SDS-PAGE and transferred to PVDF membranes (MilliporeSigma). The membranes were then blocked with 5% non-fat dry powder milk (Research Product International, IL, USA) in TBST for 1-2 hours, then incubated overnight at 4^0^C with primary antibody. In the next day, the membranes were incubated with secondary antibody and visualized their expression in iBright FL1000 Imager (Invitrogen Corporation) instrument using Chemiluminescent Substrate (Thermo Fisher Scientific, MA, USA).

### 2.9. RNA sequencing

RNA was extracted from cells using Qiagen RNeasy Mini Kit (Qiagen Inc., MD, USA). The genomic contamination from the samples was removed using DNase I treatment (Qiagen). The RNA concentration and purity were measured using a Nanodrop spectrophotometer, 1.2% agarose gel electrophoresis, and a bioanalyzer respectively. The RNA integrity value of more than 8.00 was considered to be pure in our lab and processed for sequencing. The library preparation, sequencing and analysis was performed at Novogene Corporation Inc. (California, USA) using Hiseq Illumina platform. Briefly, RNA integrity was again assed before running the sequencing, where we considered RIN >8.00 for further processing and sequencing. The high-quality RNA was fragmented and utilized for cDNA synthesis. The adaptors were ligated to the cDNA fragments to create the library and ran them into the Illumina Hi-seq platform for transcriptome analysis.

For the whole-transcriptome analysis, the raw sequencing data undergoes quality check-up and filtered to remove low-quality reads. Furthermore, they were aligned with the human reference genome (e.g., GRCh38) using HISAT2 tool. The counts were quantified using featureCounts. To identify DEGs, DESeq2 Bioconductor R-package was used. The transcriptomic data are deposited at the Gene Expression Omnibus (GEO) repository with the dataset identifier GSE304110.

### 2.10. Proteomic analysis

Cells with or without siMCM10 treatment, underwent proteomic analysis via nanoscale liquid chromatography coupled with high resolution tandem mass spectrometry (nanoLC-HR-MS/MS). Cells were lysed in 100 µL of LYSE buffer (PreOmics Inc., MA, USA), heated at 95°C for 10 minutes, and sonicated. Protein concentration was determined using the BCA Protein Assay Kit (Thermo Fisher Scientific), and 100 µg of protein was digested with the iST kit (PreOmics Inc.) using Lys-C and Trypsin at 37°C for 1 hour. The digested peptides were purified, dried, and resuspended in 100 µL of 0.1% formic acid in water which were further cleaned, and resuspended in 50 µL of 4% acetonitrile, 0.1% formic acid. Peptide concentration was assessed with the Pierce Quantitative Fluorometric Peptide Assay kit (Thermo Fisher Scientific), and samples were adjusted to 1 µg/µL before LC-MS/MS analysis.

The LC system (Dionex UltiMate™ 3000, Thermo Fisher Scientific) utilized a 50-cm reverse-phase column (bioZen 2.6 µm Peptide XB-C18) and a 2-cm RP-HPLC trap column (PepMap 100 C18). Peptides were separated with a gradient of 4–35% buffer B over 150 minutes. The peptides were ionized using a Nanospray Flex Ion Source (Thermo Fisher Scientific) and analyzed with a Q-Exactive Plus mass spectrometer, acquiring MS1 scans at 70,000 resolution and MS2 scans at 35,000 resolutions. Fragmentation was performed on the top 12 peptide ions, with dynamic exclusion set to 30 seconds.

Proteomic data were analyzed with Proteome Discoverer (PD) v2.5.0.400 (Thermo Fisher Scientific), using a 1% false discovery rate and the Human proteome database (UniProt). Analysis parameters included HCD MS/MS, up to 2 missed cleavages, and specific mass tolerances. Protein identification required at least two high-confidence peptides. Data were further processed with Scaffold Q+S for quantification and statistical analysis using the Mann-Whitney U Test, with a significance level of 0.05. The raw proteomic data are deposited to the ProteomeXchange Consortium via the PRIDE partner repository with the dataset identifier PXD066581 and 10.6019/PXD066581, and are available via ProteomeXchange with identifier PXD066581.

### 2.11. Organoid culture

The Hispanic patient-derived organoid (PDO) line “HCM-CSHL-0248-C19” was obtained from the American Type Culture Collection (ATCC) and maintained using Organoid Growth Kit 1A (ATCC: ACS-7100™), supplemented with Primocin (InvivoGen, CA, USA) to prevent microbial contamination. The growth kit includes an optimized formulation of recombinant proteins, small molecules, and supplements to support organoid viability and expansion.

Briefly, organoids were rapidly thawed in a 37°C water bath, and the cell pellet was resuspended in ice-cold growth factor-reduced Matrigel (Corning Inc.). The suspension was plated as dome structures in 6-well plates (Corning Inc.) and incubated at 37°C to allow Matrigel solidification. Each well was overlaid with 2.0 mL of complete organoid culture medium supplemented with Y-27632 (MCE, USA), a ROCK inhibitor known to enhance cell survival and proliferation. Media were refreshed every 2–3 days, and organoids were passaged approximately every 8–12 days.

### 2.12. CRISPR-mediated MCM10 Knockout (KO) in organoid

CRISPR-mediated lentiviral plasmids encoding Cas9, puromycin resistance, and MCM10-specific single guide RNAs (sgRNAs) were constructed and packaged by a commercial provider, VectorBuilder Inc. (IL, USA) to produce ready to infect lentiviral particles targeting MCM10. The viral particles were then used to transduce Hispanic PDO at an optimized multiplicity of infection (MOI) of 10. Following transduction, organoids were subjected to puromycin selection for 7–10 days to enrich for MCM10 knockout (KO) clones, which were subsequently used in downstream experiments. Two sgRNAs targeting MCM10 were tested; however, only one demonstrated efficient gene knockout in the organoid model and was selected for further functional analysis. The KO was evaluated using western blotting and considered for further experiments.

### 2.13. Organoid growth and death assay

Organoid viability and cytotoxic response following MCM10 KO were assessed using the Cyto3D™ Live-Dead Assay Kit (TheWell Bioscience, NJ, USA), according to the manufacturer’s instructions. Post-KO, organoids were seeded as Matrigel domes, with approximately 100 cells per dome, in X-well cell culture chambers (SARSTEDT, Inc., NC, USA). At both 2 and 10 days post-seeding, 2 µL of the Cyto3D reagent was added per 100 µL of culture medium, followed by incubation for 15–20 minutes at 37°C. Live cells (acridine orange-positive, green fluorescence) and dead cells (propidium iodide-positive, red fluorescence) were visualized and imaged using a Zeiss LSM 700 confocal fluorescence microscope.

### 2.14. Statistical analysis

All data are presented as the mean ± standard error of the mean (SEM) based on at least three independent experiments. Statistical significance was determined using the Student’s t-test in GraphPad Prism version 10.0 software (GraphPad Software, Inc., CA, USA) except proteomics where ue used Mann-Whitney U Test. A p-value of less than 0.05 was considered statistically significant.

## 3. RESULTS

### 3.1. SSP genes are overexpressed in CRC cell lines and human tissues

In a previously published study from our laboratory, we identified 28 SSP genes associated with CRC using microarray and RNA-seq datasets obtained from the Gene Expression Omnibus (GEO), The Cancer Genome Atlas (TCGA), and Oncomine^27^. To validate the differential expression of these SSP genes *in vitro*, we assessed their expression in six commercially available CRC cell lines (SW1116, SW480, SW620, SW48, Caco-2, and WiDr) and compared to the normal colon cell line (CCD841) using qRT-PCR. Of the 28 identified genes, the expression of three genes (*ESPL1*, *GPX2*, and *TRIB3*) was not detected in normal colon cell lines and, therefore, these genes were excluded from further analysis. Among the remaining 25 genes, 12 genes (*BCL2L1, BCL2L12, CCNB1, CDK1, CDK4, CHEK1, CSE1L, FOXM1, MCM10, NUDT1, PDCD2L,* and *PRDX4*) were significantly upregulated in all six CRC cell lines compared to normal colon cells (Fig. 1A). Of these, three cell lines (SW1116, SW480 and SW620) were KRAS positive, while the other three (SW48, Caco-2, and WiDr) were KRAS negative. However, no clear association could be established between KRAS-mediated signaling and expression of the SSP genes selected for our study, as all 12 genes were found to be significantly upregulated in all six cell lines. We further explored the transcript-level expression of these 12 SSP genes at different stages of CRC by using commercially available Colon Cancer cDNA Array IV from OriGene Technologies Inc., (Rockville, MD, USA). Ten out of the 12 genes (*BCL2L1, CCNB1, CDK1, CDK4, CHEK1, CSE1L, FOXM1, MCM10, PDCD2L,* and *PRDX4*) were found to be upregulated in the early (Stage I and II) and late (Stage III and IV) stages of CRC when compared to normal colon tissue samples (Fig. 1B). However, we could not detect any differential expression for NUDT1 and BCL2L12 across the different stages of CRC (Supplemental Fig. 1), while using the cDNA array. The overexpression of these 10 SSP genes in both early and late stages of CRC suggests their potential involvement in CRC progression from tumor initiation to advanced stages.

**Fig. 1.**
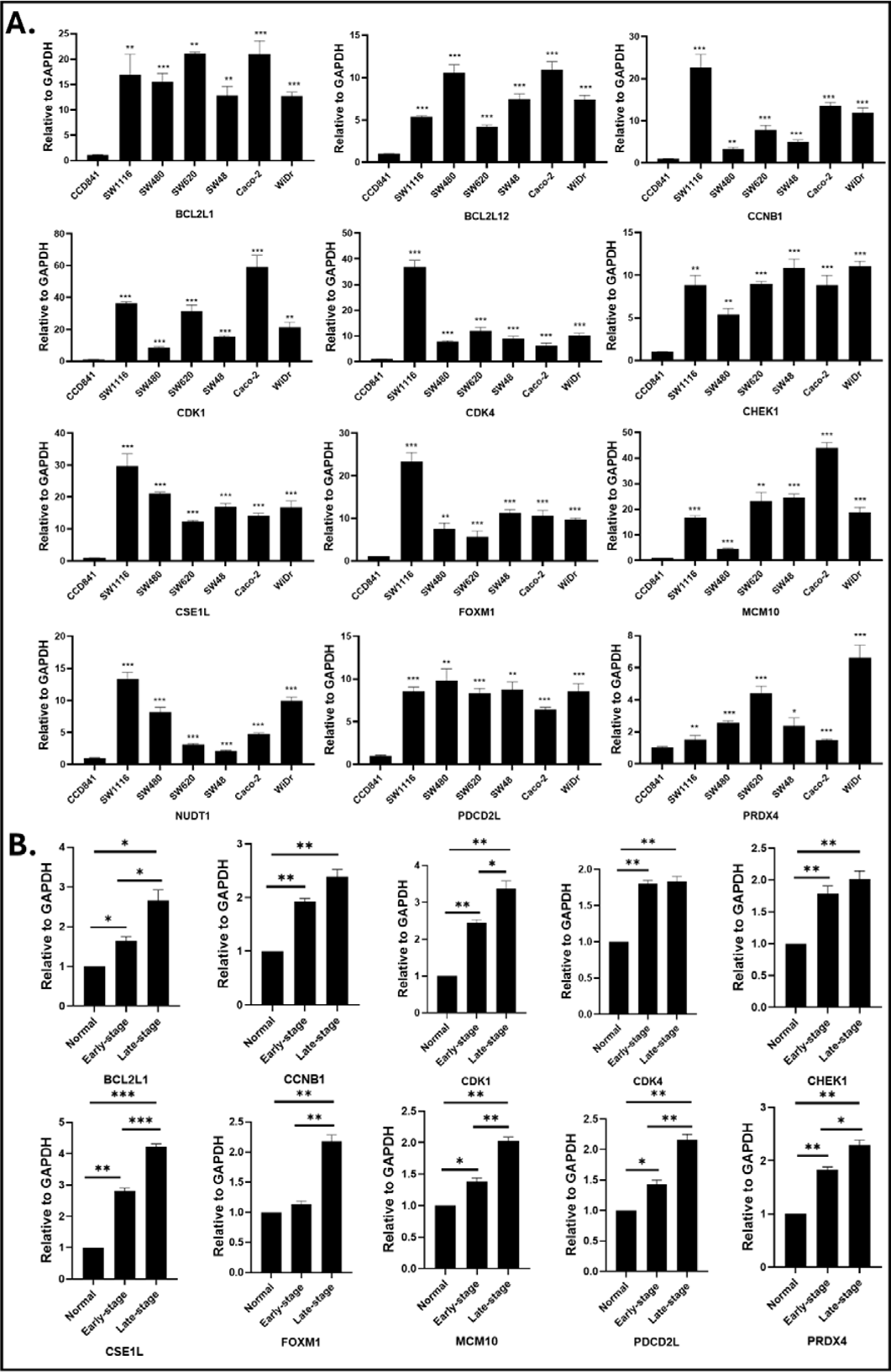
The SSP genes are significantly upregulated in CRC cells and in different stages of tumors. **A.** Expression of targeted SSP genes were quantified using qRT-PCR in six CRC cell lines compared to normal colon cells CCD841. The quantitative analysis demonstrated significant upregulation of 12 SSP genes including *BCL2L1, BCL2L12, CCNB1, CDK1, CDK4, CHEK1, CSE1L, FOXM1, MCM10, NUDT1, PDCD2L,* and *PRDX4* in six CRC cell lines, SW116, SW480, SW620, SW48, Caco-2, and WiDr, compared to normal colon cell, CCD841. **B.** The stage-specific expression of SSP genes were evaluated in both early (Stage I and II) and late stages (Stage III and IV) CRC tissues compared to the adjacent normal tissues. The significant over expressions of *BCL2L1, CCNB1, CDK1, CDK4, CHEK1, CSE1L, FOXM1, MCM10, PDCD2L,* and *PRDX4* were determined in human CRC tissues across early and late stages using cDNA array. Each bar represents relative gene expression, calculated from at least three biological replicates of this study. *(* P<0.05, ** P< 0.01, ***P<0.001*).

### 3.2. SSP proteins are differentially expressed in different stages of CRC tissue samples validating their role in CRC progression

Using human CRC TMA slides, we evaluated the protein expression levels of the ten significantly differentially expressed SSP genes in early- and late-stages CRC tissues. This analysis was performed using highly sensitive immunohistochemistry techniques that employed streptavidin-enzyme conjugates to detect biotinylated primary antibodies. Nine proteins (BCL2L1, CCNB1, CDK1, CDK4, CHEK1, CSE1L, MCM10, PDCD2L, and PRDX4) were overexpressed across all CRC tissues, irrespective of tumor pathological stage compared to their matched adjacent normal tissues (Fig. 2). In contrast, FOXM1 did not exhibit a significant difference in expression between tumor and normal tissues (Fig. 2). The consistent overexpression of these nine proteins across various CRC stages suggests effective translation of mRNA to protein, indicating their involvement in CRC progression from early development to advanced disease. Overall, these findings support the hypothesis that the upregulation of these proteins may contribute to CRC tumorigenesis.

**Fig. 2.**
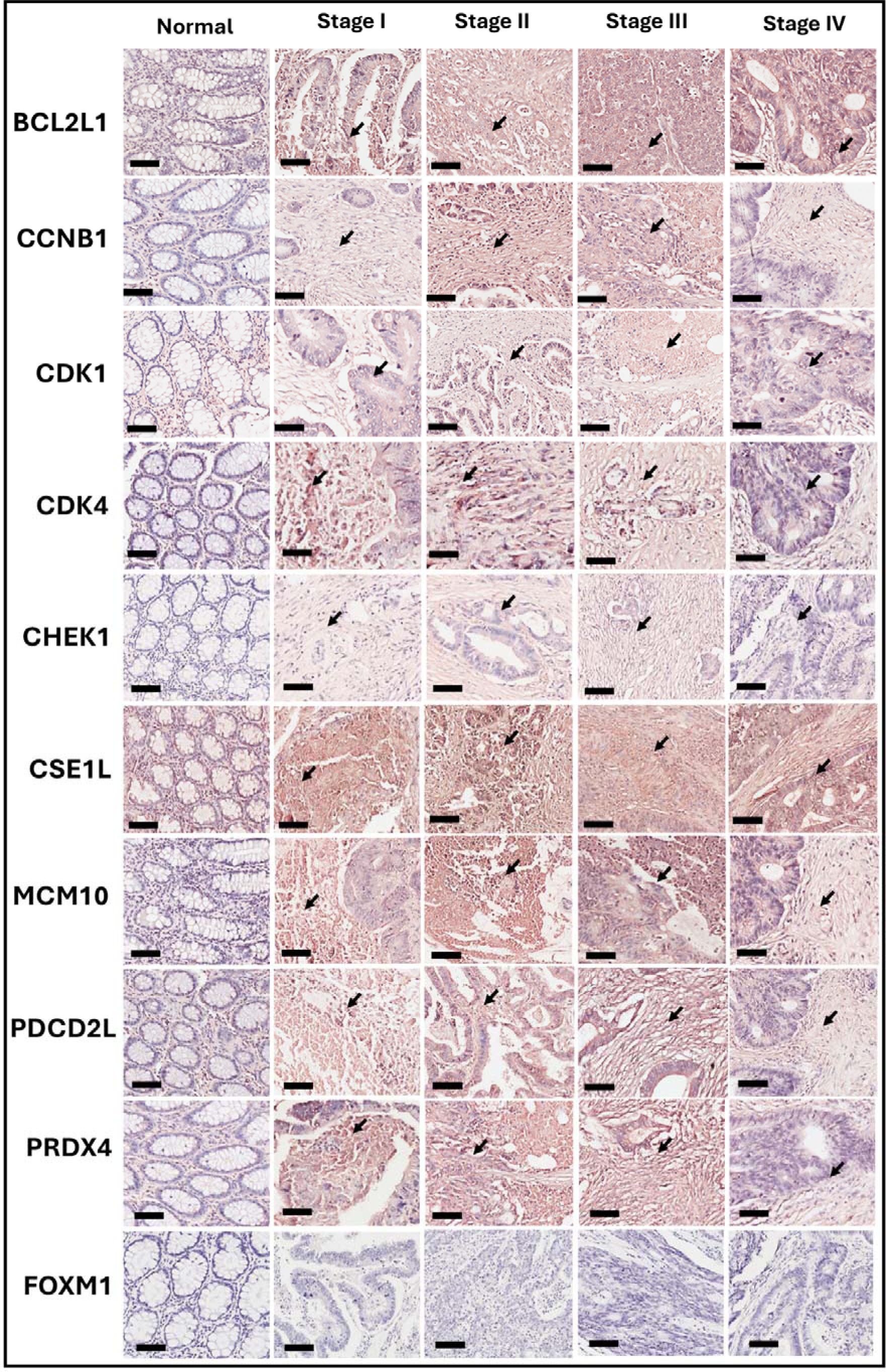
The SSP proteins are overexpressed in different stages of human CRC tissues. The significantly upregulated 10 SSP genes expression was further evaluated in protein level using Immunohistochemical assay. Nine of the proteins, including BCL2L1, CCNB1, CDK1, CDK4, CHEK1, CSE1L, MCM10, PDCD2L, and PRDX4, show upregulation at different CRC stages compared to their adjacent normal tissues. The imaging was performed with an Aperio CS2 slide scanner at 200x magnification. The staining intensity was blindly evaluated by three independent observers in a 4-tier scoring system (0=negative, 1=weak, 2=moderate, 3=strong). Positive staining is indicated by brown color (black arrow). Scale bar = 300 µm; magnification = 200x.

### 3.3. The SSP genes are differentially expressed in Hispanic CRC tissues

CRC is widely recognized for its disproportionate impact on certain racial and ethnic minority groups in the U.S^29^. Given the alarming increase in the incidence of early-onset and late-stage CRC among the Hispanic population, there is a critical need to better understand the underlying genetic factors contributing to these disparities. Emerging evidence suggests that ethnicity-specific genetic makeup, along with age- and stage-related variations, may significantly influence the expression of key genes involved in CRC pathogenesis^30^. Our results indicated that nine SSP genes with elevated transcript levels in CRC cells and tissues also exhibit high protein expressions in tumor tissues, supporting their potential significance in CRC carcinogenesis. Furthermore, we wanted to check if ethnicity-based differences might unveil their distinct molecular patterns in the Hispanic population. This led us to evaluate the expression levels of the nine SSP genes, the transcripts of which were significantly upregulated across the six CRC cell lines (SW1116, SW480, SW620, SW48, Caco-2, and WiDr) and in different stages of CRC tissues. More importantly the upregulation was consistent at the protein levels within the CRC tissues compared to normal adjacent tissue (NAT) samples. The ethnicity-based quantitative analysis was performed in CRC tissues from self-identified Hispanic (Spanish; n=18) and non-Hispanic white (NHW) patients (Russian; n=18), as well as in respective NAT samples (n=3). The qRT-PCR results revealed differential expression of all nine of the genes analyzed. Specifically, *BCL2L1, CCNB1, CDK1, CHEK1, CSE1L, MCM10, PDCD2L,* and *PRDX4* were significantly upregulated, while *CDK4* was significantly downregulated in Hispanic CRC tissue samples compared to NHWs (Fig. 3A), suggesting their potential role in Hispanic CRC disparities. In contrast, among the NAT samples, six genes exhibited significant differential expression between the two ethnic groups. However, these expression patterns differed from those observed in tumor samples (Supplemental Fig. 2). For instance, *CCNB1* and *CSE1L* were significantly downregulated in Hispanic NAT samples but were significantly upregulated in Hispanic CRC tumor samples. *CDK1, PDCD2L,* and *CHEK1* showed higher expression in Hispanic NAT samples than in the Hispanic CRC tumors, whereas *BCL2L1* exhibited greater upregulation in Hispanic CRC tumors. Notably, *MCM10, CDK4,* and *PRDX4* did not display significant difference between the Hispanic and NHW NAT samples(Supplemental Fig. 2).

**Fig. 3.**
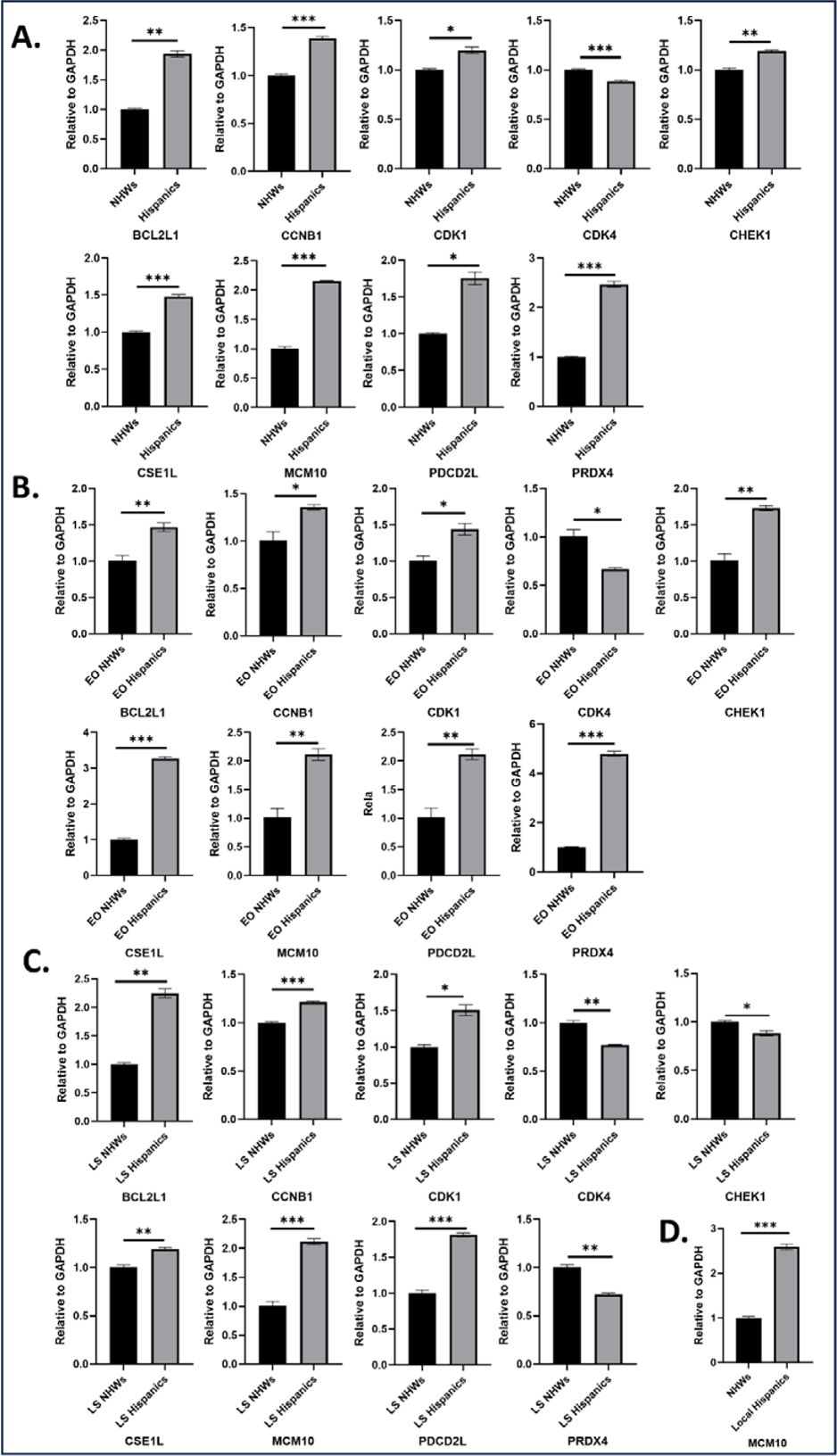
SSP genes are differentially expressed in Hispanics CRC tissues. (A). The expression of nine SSP genes including BCL2L1, CCNB1, CDK1, CDK4, CHEK1, CSE1L, MCM10, PDCD2L and PRDX4 were evaluated in Hispanic (n=18) and NHW (n=18) CRC tissues by qRT-PCR. Only CDK4 shows significant downregulation in Hispanics compared to the NHWs, whereas other eight gene shows significant upregulation among Hispanics. (**B).** The SSP genes expression was evaluated in early-onset Hispanics (n=5) and NHWs (n=5). The eight genes BCL2L1, CCNB1, CDK1, CHEK1, CSE1L, MCM10, PDCD2L, and PRDX4 shows significant upregulation in EO Hispanics compared to the EO NHW samples. CDK4 shows consistent significant downregulation among Hispanics. **(C)** The SSP gene expression was also evaluated among LS Hispanics (n=6) compared to the LS NHW (n=11) tissues. CDK4 and PRDX4 exhibited significant downregulation among Hispanics compared to the NHWs, whereas other six genes show significant upregulation. **(D)**. MCM10 expression in local Hispanics compared to the NHWs shows significant upregulation of its expression in local Hispanics which are in line with our cell lines and Hispanics derived from Spain data. Each bar represents relative gene expression, calculated from at least three biological replicates of this study. *(* P<0.05, ** P< 0.01, ***P<0.001*).

Given the rising incidence of early-onset and late-stage CRC in Hispanics, we further evaluated the expression of these nine SSP genes in CRC tissues from early-onset Hispanics (n=5) and NHWs (n=5). The findings revealed significant upregulation of *BCL2L1, CCNB1, CDK1, CHEK1, CSE1L, MCM10, PDCD2L* and *PRDX4*, and downregulation of *CDK4* in early-onset CRC (EOCRC) tissues from Hispanics compared to NHWs (Fig. 3B). These results suggest that these genes may contribute to CRC disparities in Hispanics, particularly by influencing disease progression in early-onset cases.

It is well documented that CRC disparities in Hispanics are influenced by limited access to the healthcare facilities, often leading to diagnosis at advanced stages of cancer^31^. Building on our findings from EOCRC, we hypothesized that SSP genes may also be differentially expressed in late-stage (stage III and IV) CRC (LSCRC) among Hispanics compared to NHWs. Therefore, we analyzed the expression of the 9 SSP genes in late-stage Hispanic CRC tissues (n=6) and NHWs (n=11). Our results demonstrated significant upregulation of *BCL2L1, CCNB1, CDK1, CSE1L, MCM10,* and *PDCD2L*, and downregulation of *CDK4, CHEK1,* and *PRDX4* in Hispanic CRC tissues compared to NHW samples (Fig. 3C).

In light of the consistent upregulation of *MCM10* observed in both EOCRC and LSCRC among Hispanic patients, and its lack of differential expression in NAT samples between Hispanics and NHWs, we selected this gene for further investigation within this ethnic group. To validate our findings in a representative U.S.-based Hispanic population, we examined the expression of *MCM10* in locally sourced Hispanic CRC tissues from self-identified Hispanic patients residing in Texas. This was important since our initial Hispanic CRC samples were derived from individuals in Spain, whose genetic backgrounds may differ from U.S. Hispanics. CRC tissue samples from 30 U.S.-based Hispanic patients were obtained through a commercial biorepository (iSpecimen), and qRT-PCR analyses confirmed a similar trend of *MCM10* upregulation compared to NHW samples (Fig. 3D). Collectively, these findings strengthen the hypothesis that certain SSP genes, including *MCM10*, may contribute towards the observed disparities in CRC disparities among Hispanics in the U.S. and highlight *MCM10* as potential biomarker and therapeutic target for this population.

### 3.4. MCM10 knockdown inhibits CRC cell proliferation, colony formation, invasion, and migration

MCM10, a crucial component of DNA replication, is markedly overexpressed in several cancers, including ovarian cancer, prostate cancer, gastric cancer, uterine corpus endometrial carcinoma, and breast cancer^32–37^. Despite MCM10 exploration in other cancer types, its role in CRC remains largely understudied. Given the consistent upregulation of MCM10 in early-onset and late-stage Hispanic CRC patients studied, we further investigated its oncogenic role *in vitro* and its potential contribution to CRC disparities.

Using siRNA, MCM10 gene was knocked down in Caco-2 and SW480 cell lines. In both the cell lines, >60% depletion of MCM10 expression was achieved as compared to their negative control (NC) group which contained a scrambled siRNA (Fig. 4A, B). Thereafter, the effect of MCM10 knockdown was evaluated through a series of *in vitro* functional and molecular assays. We performed CCK-8 cell proliferation assay, colony formation assay, Transwell invasion and migration assay in CRC cells pre- and post- MCM10 knockdown. The results showed that MCM10 knockdown significantly inhibited cell proliferation and growth in both Caco-2 (Fig. 4C) and SW480 (Fig. 4D) cell lines at 24, 48, and 72 hours respectively. Similarly, there was a significant reduction in the number of colonies in both cell lines post-transfection (Fig. 4E-F). Our data underscores the importance of MCM10 in promoting growth and proliferation in CRC carcinogenesis, *in vitro*.

**Fig. 4.**
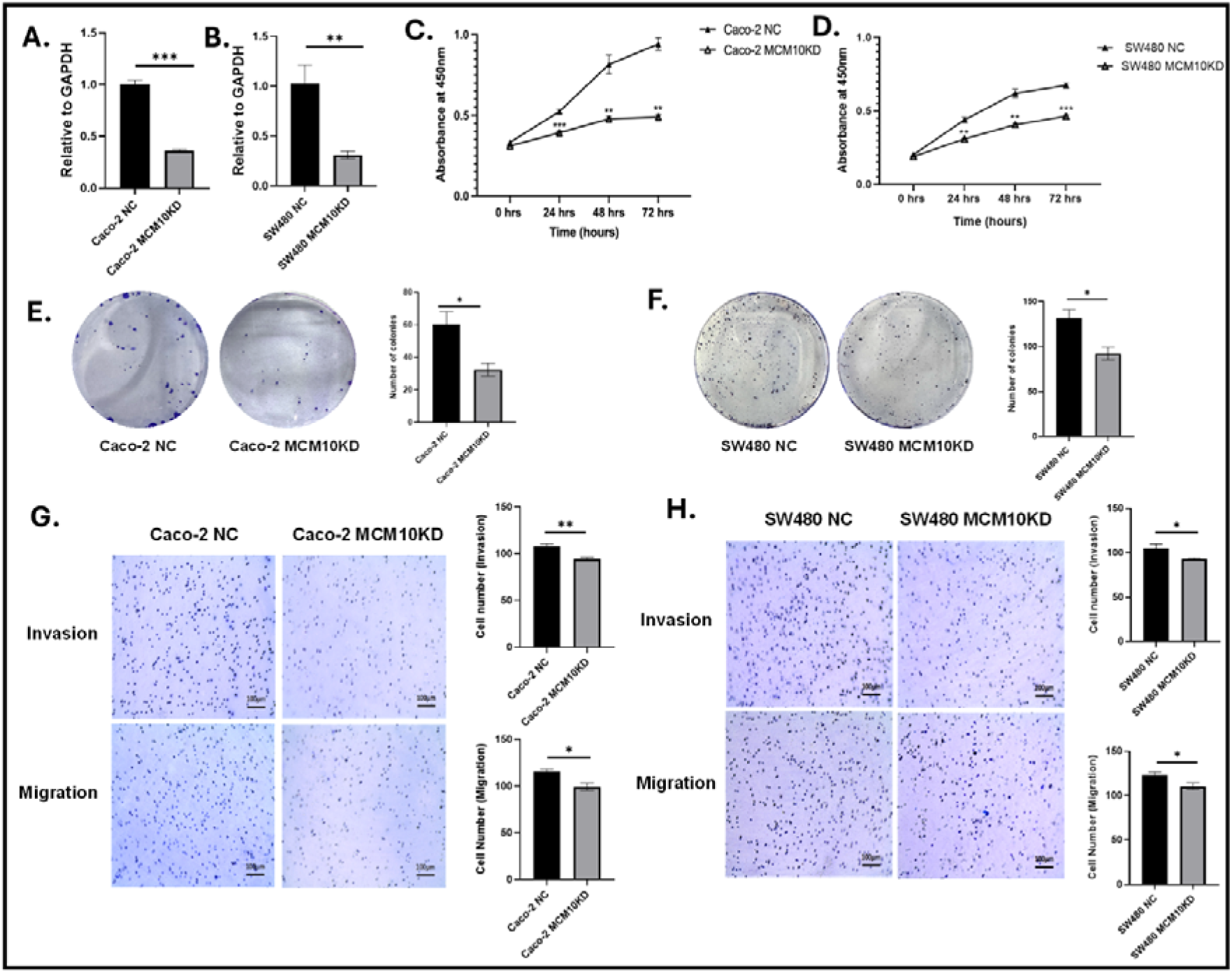
*MCM10* promotes proliferation, invasion and migration of CRC cells. (A-B). MCM10 knockdown in Caco-2 and SW480was measured by qRT-PCR. The quantitative results shown more than 60% of MCM10 knockdown in both cell lines. **(C-F)** CCK-8 mediated proliferation assay of Caco-2 and SW480 cells showed significant reduction of cell growth 24h, 48h and 72h respectively. The knockdown of MCM10 significantly reduced the number of colonies in both cell lines. The reduction of both cell proliferation and growth indicate that MCM10 could be a carcinogenic gene in CRC. **(G-H)** The MCM10 knockdown significantly reduced the cellular invasiveness and migratory characteristics of the cells compared to their NC group. These data indicate that MCM10 could be a potential target reducing the CRC carcinogenesis. The data was shown as meanLJ±LJSEM compared to their negative control group. (*\* P<0.05, ** P< 0.01, ***P<0.001*). Scale bar= 300 µm.

Next, we evaluated the invasive and migratory characteristics using both cell lines; the results showed that the knockdown of MCM10 significantly reduced the cellular invasiveness and migration in Caco-2 and SW480 cell lines (Fig. 4G-H). These findings suggest that MCM10 plays a crucial role in facilitating cell invasion and migration in CRC and could also be associated with tumor progression.

### 3.5. MCM10 knockdown enhances apoptosis, ROS production, and cell-cycle arrest in Caco-2 and SW480 cells

To evaluate apoptosis and cell-cycle arrest, flow cytometric analysis was performed in both the cell lines after MCM10 knockdown. In this study, MCM10 knockdown displayed a significant reduction in the proportion of cells in the G1/G0 phase and an increase in the cells in G2/M phase. The accumulation of cells in G2/M phase suggests a disruption in the normal cell cycle progression, with cells failing to transition from G2 to Mitosis. This indicates that MCM10 knockdown reduced cell growth by inducing cell cycle arrest at G2/M phase, potentially suppressing carcinogenesis (Fig. 5A-B). In addition, the annexin V-FITC apoptosis assay revealed that knockdown of MCM10 significantly enhanced the cellular apoptosis in both cell lines as compared to their negative control (Fig. 5C-D). Therefore, the results from both cell cycle and apoptosis assays suggested that MCM10 knockdown may reduce tumorigenesis by activating apoptotic cell death and cell cycle arrest.

**Fig. 5.**
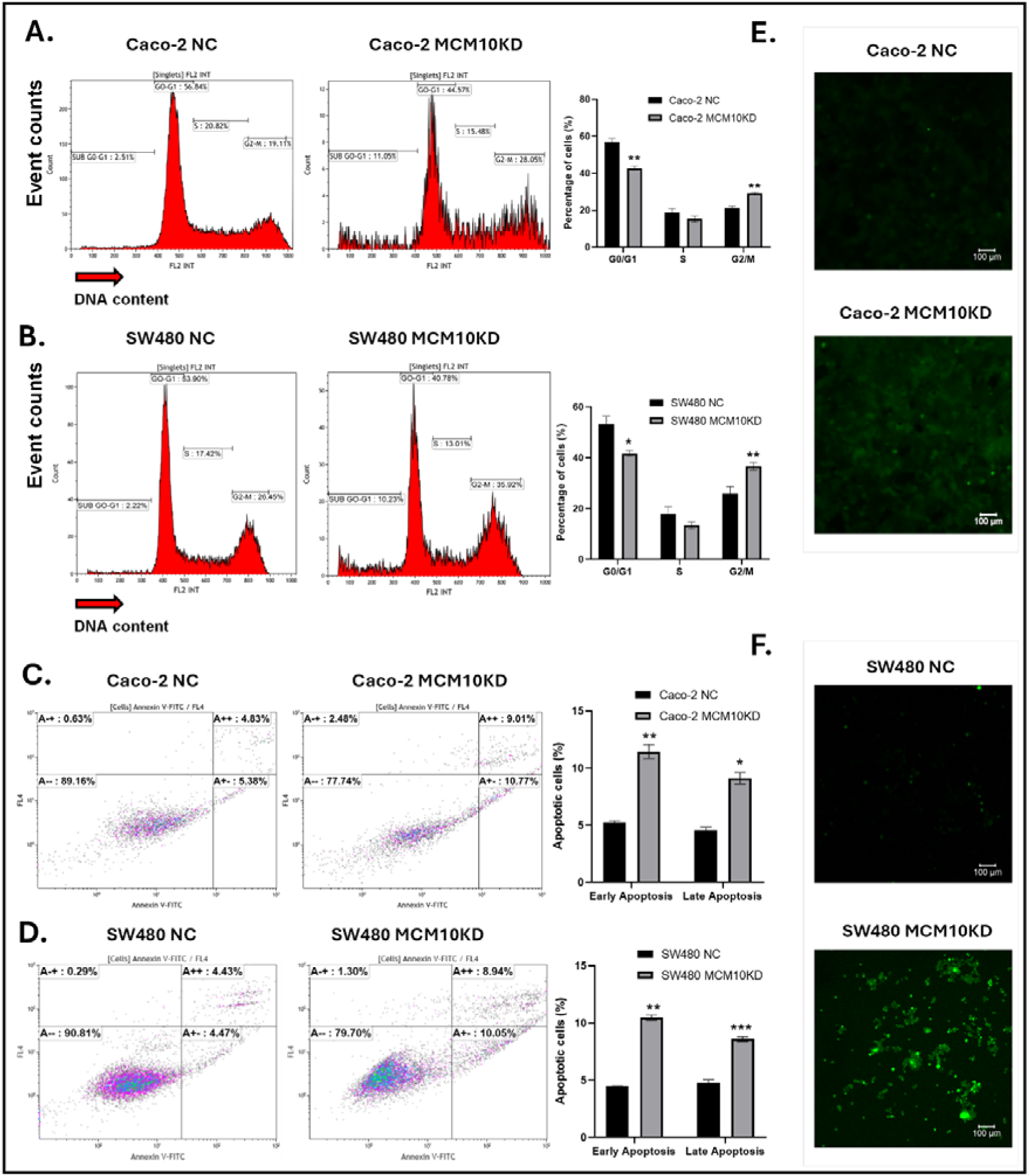
MCM10 knockdown induced cellular apoptosis, cell cycle arrest and ROS production in CRC cells. (**A-B).** The flow cytometric analysis was performed to evaluate the cell cycle arrest after MCM10 knockdown in both Caco-2 and SW480 cells. The depletion of MCM10 significantly reduced the number of cells at G1/G0 and increased at G2/M phase respectively. **(C-D).** Likewise, the apoptosis level of both cell lines was determined by flow cytometric analysis using annexin V-FITC. The result demonstrated that MCM10 knockdown significantly increased the early and late apoptosis in Caco-2 and SW480 cell lines compared to their respective NC group. (**E-F).** The ROS was evaluated to check whether cell cycle arrest and apoptosis process was induced by reactive oxygen or not. The knockdown of MCM10 in Caco-2 and SW480 cells potentially activate the cellular ROS production to promote cell cycle arrest and apoptosis pathway to reduce the carcinogenesis compared to their respective NC group. The results are representative of at least three independent experiments. (*\* P<0.05, ** P< 0.01, ***P<0.001*). Scale bar= 100µm.

To further elucidate the mechanism underlying cellular death after MCM10 knockdown, we performed ROS assay. The MCM10 depleted cells dramatically increased ROS production in Caco-2 and SW480 cells as compared to NC group (Fig. 5E-F). This finding indicates the induction of ROS may induce apoptosis and growth arrest *in vitro* after MCM10 knockdown in both cell lines.

### 3.6. An insight into the molecular mechanism underlying reduced CRC progression after MCM10 knockdown and the potential role of PPFIA1

To decipher the downstream signaling underlying reduced CRC carcinogenesis as a result of MCM10 knockdown, we conducted high throughput RNA-sequencing (RS) in both Caco-2 and SW480 cell lines with and without MCM10 knockdown. We identified 26 and 44 differentially expressed genes (DEGs) in Caco-2 and SW480 cell lines with MCM10 knockdown, respectively, with a log2 fold change cutoff of >0.95 (>1.9 fold) and p-adjusted value (with FDR correction) of < 0.05 (Fig. 6A-B). Among these, five DEGs were common to both cell lines (Supplemental Fig. 3), including MCM10, which was downregulated, confirming a successful knockdown. The remaining four DEGs include ZNF268, ENTPD8, PPFIA1, and ZNF33B could be regulated by MCM10 mediated CRC progression.

**Fig. 6.**
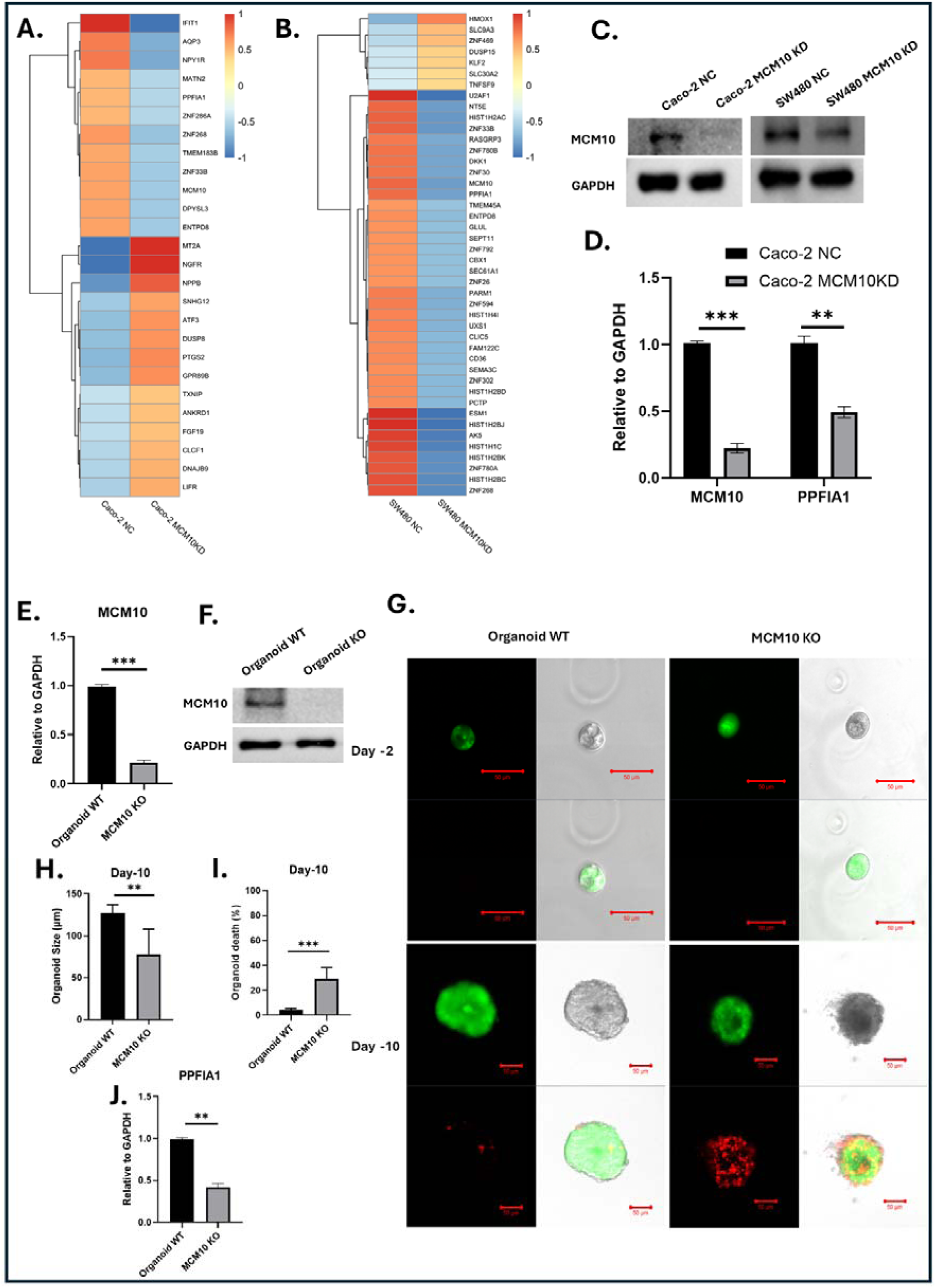
**Transcriptomic/proteomic analysis post *MCM10* Knockdown in CRC cells and validation of findings in Hispanic organoid. (A-B)**. The downstream signaling was evaluated by performing high-throughput RS and high-resolution proteomic- analysis, respectively. The heatmap from RS showed differentially expressed genes from Caco-2 and SW480 cells post-MCM10 knockdown. Overall, 26 and 44 DEGs were identified in Caco-2 and SW480 cell lines with MCM10 knockdown, with a log2 fold change cutoff of >0.95 and p-adjusted value (with FDR correction) of < 0.05. **(C).** Protein expression of MCM10 after transfection which was evaluated by western blotting where it shows clear downregulation of its expression in MCM10-KD group. **(D).** The qRT-PCR was performed to evaluate the expression of PPFIA1 as key differentially expressed genes and protein in response to MCM10 KD where it significantly downregulated along with MCM10 in transfected group compared to the NC group. A significant downregulation of PPFIA1 was observed along with MCM10 after the transfection **(E-F).** The MCM10 KO efficiency was measured by qRT-PCR in transcript level and western blotting in protein level. **(G-I)** Furthermore, the size of organoid was evaluated after MCM10 KO. The MCM10 KO significantly reduced the organoid size compared to the WT group. Similarly, the live (green fluorescence by premixed acridine orange dye) and dead (red fluorescence by propidium iodide dye) assay revealed that MCM10 KO significantly induces the apoptosis in order to reduce the carcinogenesis. **(J)** Notably, MCM10 KO significantly reduced the expression of PPFIA1 in Hispanic organoid model which are in line with our cell lines demonstration. The data was shown as meanLJ±LJSEM compared to their negative control group and representative of at least three independent experiments. (*\* P<0.05, ** P< 0.01, ***P<0.001*). Scale bar= 100µm.

To further identify the potential downstream effects of MCM10 knockdown, we evaluated the translational changes in Caco-2 cells by performing Mass Spectrometry (MS) based proteomic analysis. We also performed western blotting in MCM10 depleted cells and confirmed downregulation of MCM10 expression in both Caco-2 and SW480 cell lines (Fig. 6C). The MS analysis revealed 78 differentially expressed proteins in Caco-2 cells after MCM10 knockdown as compared to the control group (Supplementary Table 4). The integration of RS and MS data revealed PPFIA1 as the only differentially expressed transcript and protein in response to MCM10 knockdown, suggesting its potential involvement in CRC progression.

Furthermore, to validate our integrated RS (from Caco-2 and SW480 cells) and MS (from Caco-2 cells) findings indicating that PPFIA1 is consistently downregulated in response to MCM10 knockdown, we performed qRT-PCR to study the PPFIA1 expression in MCM10 depleted Caco-2 cells compared to the control group. The results demonstrated a significant reduction of PPFIA1 in the MCM10 depleted Caco-2 cells (Fig. 6D). We performed survival analysis for both MCM10 and PPFIA1, using the KM Plotter webtool, that uses data from a clinical subsystem which includes data from 3.55 million patients. We observed that elevated expression levels of MCM10 and PPFIA1 are significantly associated with reduced overall survival in CRC patients across all stages (Supplemental Fig. 4 A-B). In late-stage CRC patients, higher expression of MCM10 and PPFIA1 is also significantly correlated with poorer overall survival (Supplemental Fig. 4 C-D) These findings underscore the critical role of PPFIA1 in MCM10-mediated tumor progression and suggest that targeting the MCM10/PPFIA1 axis may represent a novel therapeutic strategy for CRC.

To address Hispanic CRC disparities, we validated our *in vitro* findings in Hispanic patient-derived CRC organoids and performed CRISPR-mediated MCM10 KO experiments. The depletion of MCM10 was confirmed in transcript level by qRT-PCR and protein level by western blotting (Fig. 6E-F).

Furthermore, to evaluate the oncogenic role of MCM10 in Hispanic context, we performed live and dead assays after MCM10 KO in an organoid model. Two days after MCM10 KO, we did not observe any changes in growth whereas the growth data after ten days revealed a significant reduction of organoid size in MCM10 KO group compared to the control or wild type (WT) group (Fig. 6G-H). Similar to the growth assay, we did not find any cell death 2 days post MCM10 KO in the organoid. However, a significant amount of cell death was observed in MCM10 KO organoid after 10 days of the study, which are in agreement with our cell line knockdown results of MCM10 (Fig. 6G, I). This corroborates the contribution of the SSP gene MCM10 to Hispanic CRC disparities.

Considering the functional similarities between cell lines and organoid results after MCM10 suppression, we wanted to check the potential role PPFIA1 in Hispanic CRC disparities. Therefore, we evaluated its expression in organoid model after MCM10 KO. The results were congruent with our previous results; the expression of PPFIA1 was significantly reduced in the MCM10 KO group from the Hispanic PDO model compared to the WT group (Fig. 6J). The overall study findings highlight the critical role of the MCM10/PPFIA1 axis in Hispanic CRC disparities and suggest that targeting this pathway may offer a novel therapeutic strategy for improved CRC management.

## 4. DISCUSSION

CRC is one of the most prevalent gastrointestinal malignancies worldwide and is the second leading cause of cancer death in the U.S^38^. The significant incidence and mortality rates associated with CRC are largely driven by complex molecular mechanisms, such as genetic mutations, epigenetic alterations, and dysregulation of signalling pathways that regulate cell proliferation, invasion, migration, cell cycle regulation and apoptosis^28,39^. Therefore, better understanding of these complex processes associated with CRC growth and progression is crucial to identify effective biomarker candidates and novel therapeutic targets, which could lead to effective management and improved outcome of the diseases.

The significance of this study is two-fold - i) it identifies stage-specific SSP genes through comprehensive screening of CRC clinical samples, TMAs, cDNA arrays and cell lines; and ii) it investigates the role of the SSP gene MCM10 in contributing to CRC disparities among Hispanic populations with an emphasis of early-onset and late-stage CRC.

Our integrated transcript and protein level validations revealed significant upregulation of 9 out of 28 SSP genes, BCL2L1, CCNB1, CDK1, CDK4, CHEK1, CSE1L, MCM10, PDCD2L, and PRDX4 across various stages of CRC, underscoring their potential as biomarkers or therapeutic targets. These genes are known to be involved in cell cycle progression, apoptosis resistance, DNA repair, and oxidative stress regulation, suggesting their potential as both diagnostic biomarkers and actionable therapeutic targets. Among the differentially expressed genes, BCL2L1 and PRDX4 are pivotal regulators of apoptosis and oxidative stress^40,41^ and their upregulation has been associated with tumor survival and progression by inducing oxidative stress and impairing the DNA-damage repairing process^40–42^. Similarly, overexpression of CDK1, CDK4, and CCNB1 contributes to CRC proliferation and chemoresistance via modulation of the p53 pathway and Cdc37 phosphorylation^43,44^. Genes like PDCD2L and CSE1L enhance tumor aggressiveness by promoting G2/M transition and inhibiting apoptotic signaling^45–47^. Notably, CHEK1 and MCM10 are central to the DNA damage response (DDR), enabling tumor cells to evade apoptosis and maintain genomic instability—a hallmark of cance^48,49^.

CRC also disproportionately affects racial and ethnic minorities, including Hispanic populations, who often present with early-onset and advanced-stage disease^29^. Several studies have attributed these disparities to biological heterogeneity, socioeconomic barriers, and unequal access to care^50–52^. Our findings add a crucial biological dimension by revealing differential expression of several SSP genes—most notably MCM10—between Hispanics and NHWs. This suggests ethnicity-specific molecular drivers that may underlie the observed clinical disparities.

Our study identifies MCM10 as a particularly promising ethnicity-specific biomarker for CRC disparities in Hispanics. MCM10, a DNA replication licensing factor, is known to facilitate cell cycle progression and genome duplication under stress. Previous reports have linked its overexpression to poor prognosis in endometrial and lung cancers, likely through its interaction with the CCND1 axis and DDR pathways^33,35,36,53^. We extend these findings to CRC, showing for the first time that MCM10 is significantly overexpressed in early-onset and late-stage Hispanic CRC tissues, cell lines, and a patient-derived organoid model and a novel MCM10–PPFIA1 axis drives the function. Functional studies demonstrate that MCM10 knockdown in CRC cell lines (Caco-2 and SW480) reduces proliferation, migration, and invasion by arresting cells in the G2 phase and enhancing apoptosis via increased ROS production. Mechanistically, we identified PPFIA1 as a downstream effector of MCM10, suggesting that MCM10–PPFIA1 axis promotes CRC progression. Importantly, MCM10 silencing in a Hispanic patient-derived CRC organoid significantly reduced organoid viability and growth, further validating its role as a potential therapeutic target.

Despite these compelling findings, a key limitation of our study is that the Hispanic-specific functional validation of MCM10 was performed using only one patient-derived organoid model. To enhance reproducibility and translation to clinical practice, future studies should expand to include a larger, ethnically diverse cohort and multiple Hispanic-derived organoid models. Furthermore, longitudinal clinical correlation and functional in vivo validation are warranted to fully establish the biomarker utility and therapeutic potential of MCM10 in CRC. Another limitation lies in the use of multiomics analysis performed on only two cell lines—Caco-2 (KRAS mutation-positive) and SW480 (KRAS mutation-negative). The rationale for this experimental design was to assess functional differences between cells with high and low MCM10 expression, as MCM10 is expressed 8–10 times higher in Caco-2 cells compared to SW480 (Fig. 1A). However, a potential weakness is the lack of observed correlation between SSP gene expression and KRAS mutation status in these cell lines. Notably, a recent study reports that Hispanic CRC patients exhibit a higher frequency of KRAS mutations but fewer BRAF and KIT mutations compared to NHW patients^54^. Although our selected cell lines differed in KRAS mutation status, they were derived primarily from individuals of White ethnicity. Therefore, direct comparisons using these models may not accurately reflect the genetic disparities between Hispanic and NHW populations. To address this gap, future studies should employ organoids derived from both Hispanic and NHW patients with known KRAS mutation status to better capture the ethnic and molecular heterogeneity in CRC.

## 5. CONCLUSION

We have used a set of 42 commercially available patient tissue samples (36 CRC and 6 NATs from Spain and Russia) to check the expression of the 9 SSP genes and confirmed the expression of MCM10 in 30 local (from Texas) Hispanic CRC samples. We have also used a Hispanic CRC PDO to validate the role of MCM10 and the expression of a downstream gene PPFIA1. Use of Hispanic PDOs is not common and commercially available Hispanic CRC PDOs are rare. Further investigation could promote the use of these genes as potential diagnostic biomarkers and therapeutic targets for Hispanics and NHW CRC patients. This study provides a novel insight into the role of several SSP genes in cancer progression and in Hispanic CRC disparities and validates the role of MCM10 in CRC with an emphasis on Hispanic EOCRC and LSCRC, by using multi-omics, molecular and cellular analyses. Thereby, establishing it as a particularly promising ethnicity-specific biomarker for CRC disparities in Hispanics.

## List of abbreviations

Abbreviation Definition

CRC Colorectal cancer

SSP Stress-survival pathway

NHW Non-Hispanic White

U.S. United States

ROS Reactive oxygen species

MCM10 Minichromosome maintenance 10

NAT Normal-adjacent tissue

siRNA Small interfering RNA

7-AAD 7-amino-actinomycin

FFPE Formalin fixed paraffin embedded

TMA Tissue microarray

PDO Patient-derived organoid

ATCC American type culture collection

sgRNA Single guide-RNA

KD Knockdown

SEM Standard error of the mean

GEO Gene expression omnibus

TCGA The cancer genome atlas

NC Negative control

RS RNA-sequencing

DEG Differentially expressed gene

MS Mass spectrometry

WT Wild type

EO Early-onset

LS DDR

Late stage

DNA damage response

## Ethics approval and consent to participate

Human tissue samples were procured from reputable international commercial vendors. Informed consent was obtained for all specimens by the vendors, with vendors affirming that collection adhered to ethical standards and complied with all relevant local and international regulations. Written consent covered broad research use, though not specifically for this study.

## Consent for publication

Not applicable.

## Availability of data and materials

The data used to generate the presented figures are published with this paper. Raw data will be made available upon request to the corresponding author. The sequencing data that support the findings of the study have been submitted to following publicly available repositories.

1. We submitted the transcriptomic data the NCBI GEO repository. The accession number is “GSE304110” and
2. The raw proteomic data are deposited to the ProteomeXchange Consortium via the PRIDE partner repository with the dataset identifier PXD066581 and 10.6019/PXD066581, and are available via ProteomeXchange with identifier PXD066581.

## Authors’ contributions

SR conceived, designed and procured funding for the project. TAH and AB provided critical inputs toward the project design. ZIK, UB, and SN conducted the experiments, analysed data, and wrote the manuscript. ZIK, UB, and SN are equally contributed to this work. AK, FAR, and AT conducted some experiments. AB analyzed and interpreted the immunohistochemistry data, reviewed and edited the manuscript. SR and TAH analyzed and interpreted the results, reviewed and edited the manuscript for publication. ICA analyzed the proteomics data reviewed and edited the manuscript. All the authors read, approved and finalized the manuscript.

## Acknowledgements

We thank Prof. Marc Cox (PI, CPRIT grant # RP210153), and Prof. Robert A. Kirken (PI, NIH/NIMHD grant # U54MD007592), University of Texas at El Paso, for their support. We are also grateful to the Biomolecule Analysis and Omics Unit (NIH/NIMHD grant # U54MD007592, to R.A. Kirken) for the proteomic analysis. We would also like to thank Prof. Michael Schatz (PI-OT2OD034190), Johns Hopkins University, for his support.

