## Supplementary material for "Stress-Survival Pathway Profiling Reveals MCM10 as a Candidate Biomarker of Hispanic Colorectal Cancer Disparities": 20251113_Supplementary file.docx

**Supplementary Figures and Tables**

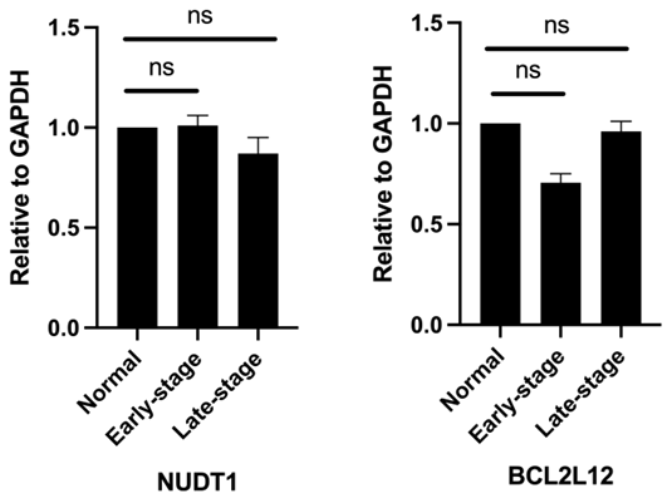

**Supplemental Fig. 1:** The qRT-PCR evaluation of NUDT1 and BCL2L12 across different stages of CRC samples using cDNA tissue micro-array, we did not find any significant differential expression of NUDT1 and BCL2L12 across different stages of CRC samples. Each bar represents relative gene expression, calculated from at least three biological replicates of this study. *(* P<0.05, ** P< 0.01, ***P<0.001*).

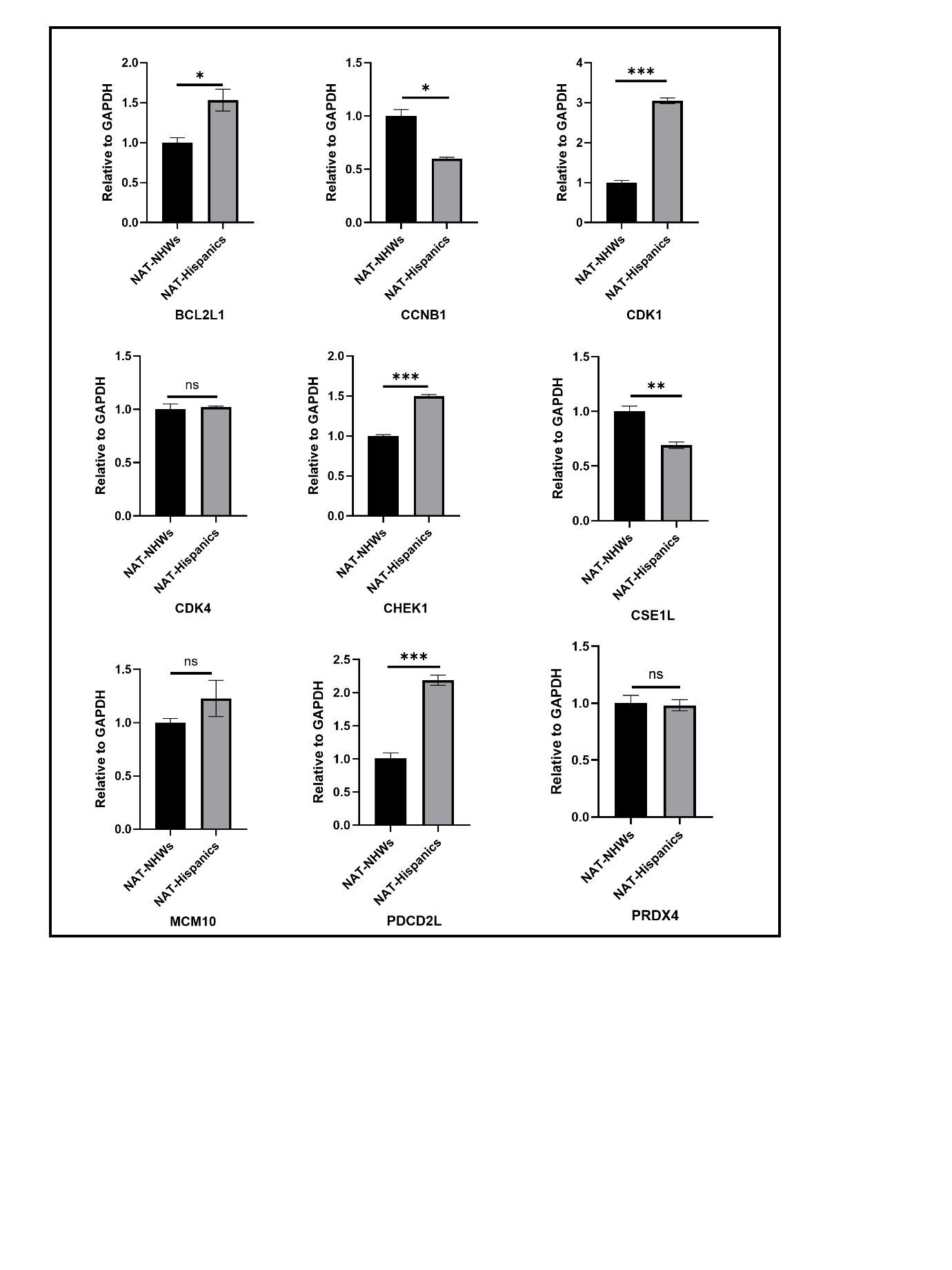

**Supplemental Fig. 2:** The qRT-PCR evaluation of NAT samples from both Hispanic (n=3) and NHW (n=3) population. The BCL2L1, CCNB1, CDK1, CHEK1, CSE1L, and PDCD2L shows significant differential expression between ethnic comparison. Whereas, CDK4, MCM10 and PRDX4 did not show significant expression changes among the groupEach bar represents relative gene expression, calculated from at least three biological replicates of this study. (* P<0.05, ** P< 0.01, ***P<0.001).

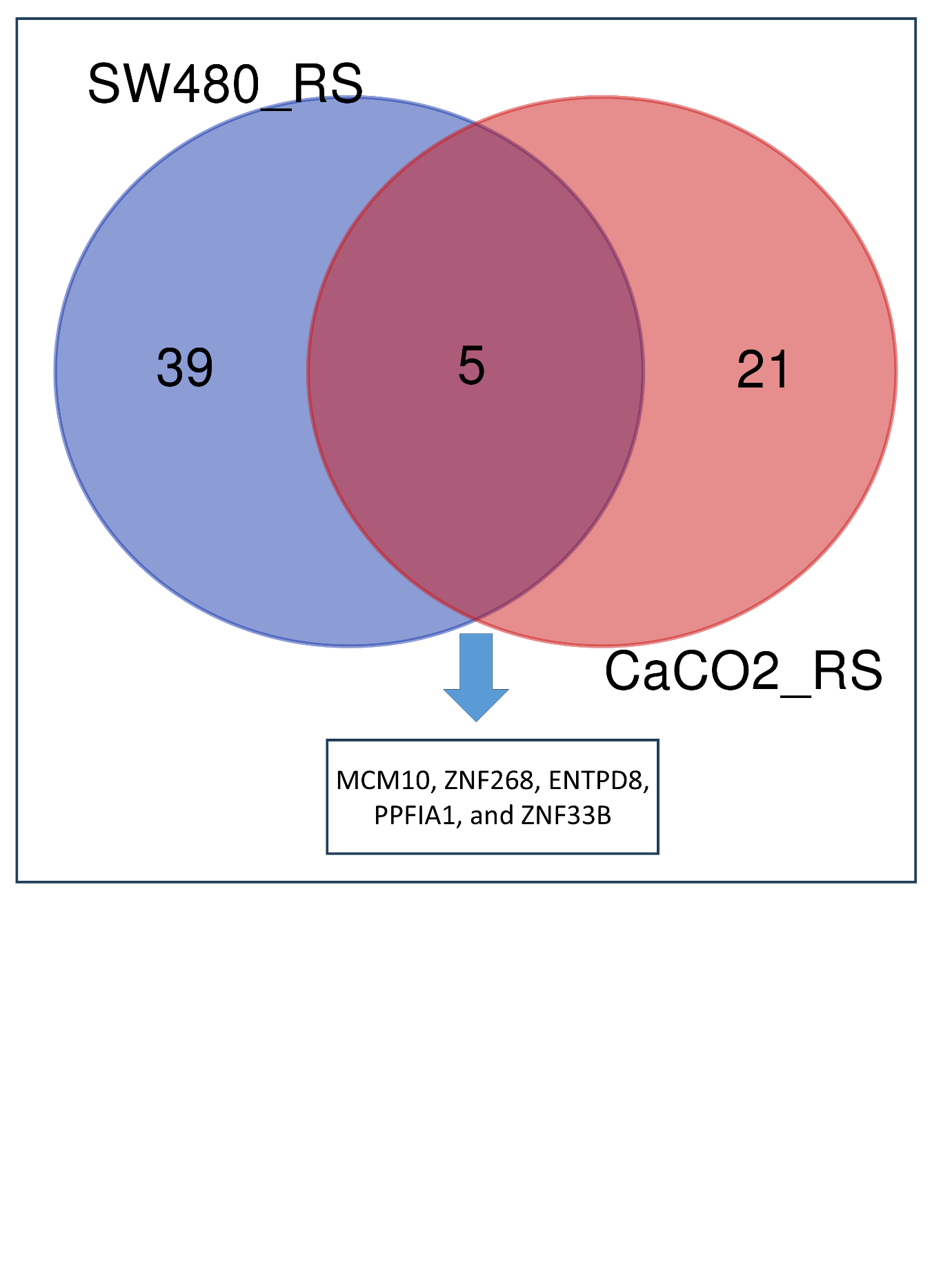

**Supplemental Fig. 3**: The differential expressed common genes from SW480 and CaCo-2 cell lines after MCM10 knockdown. The values are made with a log2 fold change cutoff of >0.95 and p-adjusted value (with FDR correction) of < 0.05.

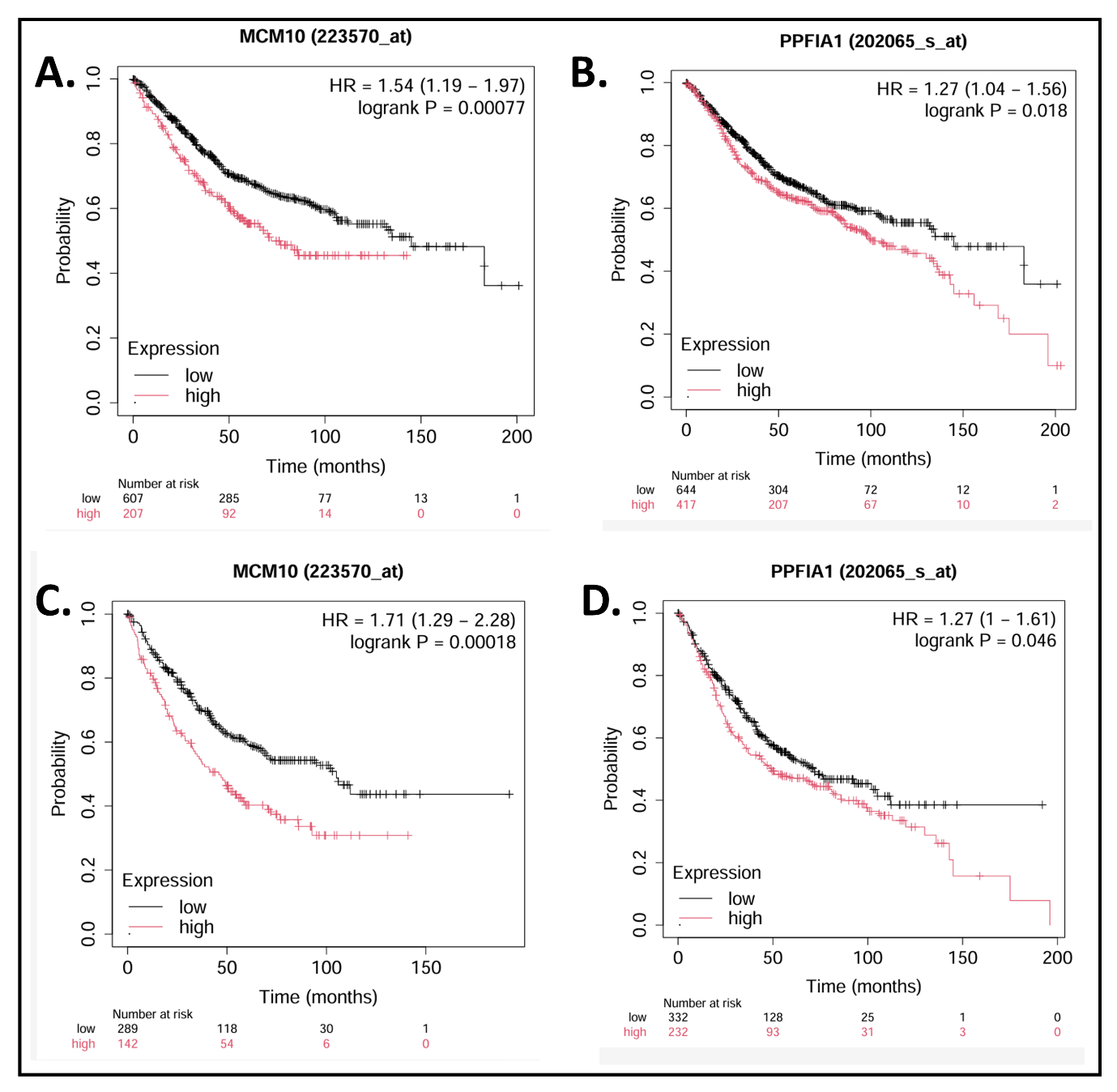

**Supplemental Fig. 4: (A-B)** Elevated expression levels of MCM10 and PPFIA1 are significantly associated with reduced overall survival in CRC patients across all stages. **(C-D)** In late-stage CRC patients, higher expression of MCM10 and PPFIA1 is also significantly correlated with poorer overall survival, potentially reflecting increased treatment failure and metastatic risk. Plots were generated using an online platform “Kaplan-Meier Plotter” using their default settings.

**Supplementary Table 1:** Patient details for both Hispanic and Non-Hispanic Whites

| **Hispanic Patients Details** | | | | | |
| --- | --- | --- | --- | --- | --- |
| Lab ID | Category | Gender | Age | Cancer Stage | Ethnicity |
| H4 | Tumour | F | 72 | IIIC | Hispanic or Latino |
| H5 | Tumour | F | 50 | IIIB | Hispanic or Latino |
| H8 | Tumour | F | 67 | Unknown | HISPANIC/LATINO |
| H9 | Tumour | F | 72 | STAGE I | HISPANIC/LATINO |
| H10 | Tumour | M | 45 | STAGE III | HISPANIC/LATINO |
| H11 | Tumour | F | 58 | STAGE II | HISPANIC/LATINO |
| H12 | Tumour | M | 50 | STAGE II | HISPANIC/LATINO |
| H13 | Tumour | M | 82 | Unknown | HISPANIC/LATINO |
| H14 | Tumour | F | Unknown | STAGE IIB | Hispanic |
| H15 | Tumour | F | Unknown | STAGE II | Hispanic |
| H16 | Tumour | F | 89 | Unknown | HISPANIC/LATINO |
| H18 | Tumour | M | 49 | STAGE IIB | HISPANIC/LATINO |
| H21 | Tumour | F | Unknown | Unknown | HISPANIC |
| H23 | Tumour | F | 59 | IIA | HISPANIC |
| H24 | Tumour | F | 64 | IIA | HISPANIC |
| H25 | Tumour | F | 88 | IV | HISPANIC |
| H26 | Tumour | M | 51 | IV | HISPANIC |
| H22 | Tumour | M | 47 | IVA | HISPANIC |
| NAT-H10 | Normal | Unknown | Unknown | Unknown | Hispanic |
| NAT-H12 | Normal | Unknown | Unknown | Unknown | Hispanic |
| NAT-H16 | Normal | Unknown | Unknown | Unknown | Hispanic |
| **Non-Hispanic White Patients Details** | | | | | |
| Lab ID | Category | Gender | Age | Cancer Stage | Ethnicity |
| NHW1 | Tumour | F | 54 | IIIC | Caucasian |
| NHW2 | Tumour | M | 57 | IIIB | Caucasian |
| NHW3 | Tumour | M | 61 | IIIA | Caucasian |
| NHW4 | Tumour | F | 62 | IIIC | Caucasian |
| NHW8 | Tumour | M | 39 | STAGE II | WHITE/CAUCASIAN |
| NHW6 | Tumour | F | 54 | IIIA | White or Caucasian |
| NHW9 | Tumour | F | 78 | STAGE IV | WHITE/CAUCASIAN |
| NHW10 | Tumour | F | 52 | STAGE IV | WHITE/CAUCASIAN |
| NHW11 | Tumour | M | 46 | STAGE I | WHITE/CAUCASIAN |
| NHW12 | Tumour | M | 52 | STAGE IV | WHITE/CAUCASIAN |
| NHW13 | Tumour | F | 73 | STAGE III | WHITE/CAUCASIAN |
| NHW14 | Tumour | F | 52 | STAGE I | WHITE/CAUCASIAN |
| NHW15 | Tumour | F | 68 | STAGE II | WHITE/CAUCASIAN |
| NHW16 | Tumour | M | 61 | STAGE III | WHITE/CAUCASIAN |
| NHW17 | Tumour | F | 54 | STAGE IV | WHITE/CAUCASIAN |
| NHW19 | Tumour | F | 46 | IIA | Caucasian |
| NHW20 | Tumour | F | 47 | I | Caucasian |
| NHW21 | Tumour | F | 48 | IIA | Caucasian |
| NAT-NHW12 | Normal | Unknown | Unknown | Unknown | Caucasian |
| NAT-NHW15 | Normal | Unknown | Unknown | Unknown | Caucasian |
| NAT-NHW17 | Normal | Unknown | Unknown | Unknown | Caucasian |
| **U.S. based Hispanic tissue details** | | | | | |
| LH1 | Tumour | M | 59 | IIA | Hispanic |
| LH2 | Tumour | M | 56 | IIA | Hispanic |
| LH3 | Tumour | M | 68 | IIA | Hispanic |
| LH4 | Tumour | F | 72 | IIA | Hispanic |
| LH5 | Tumour | M | 68 | IIA | Hispanic |
| LH6 | Tumour | F | 74 | IIA | Hispanic |
| LH7 | Tumour | M | 71 | IIA | Hispanic |
| LH8 | Tumour | M | 65 | I | Hispanic |
| LH9 | Tumour | F | 72 | IIA | Hispanic |
| LH10 | Tumour | M | 66 | IIA | Hispanic |
| LH11 | Tumour | F | 85 | I | Hispanic |
| LH12 | Tumour | F | 41 | I | Hispanic |
| LH13 | Tumour | F | 58 | IIA | Hispanic |
| LH14 | Tumour | M | 66 | I | Hispanic |
| LH15 | Tumour | F | 76 | I | Hispanic |
| LH16 | Tumour | M | 58 | I | Hispanic |
| LH17 | Tumour | M | 57 | IIA | Hispanic |
| LH18 | Tumour | M | 67 | IIA | Hispanic |
| LH19 | Tumour | F | 64 | IIA | Hispanic |
| LH20 | Tumour | M | 68 | IIA | Hispanic |
| LH21 | Tumour | M | 67 | I | Hispanic |
| LH22 | Tumour | F | 23 | I | Hispanic |
| LH23 | Tumour | F | 41 | IIA | Hispanic |
| LH24 | Tumour | F | 47 | IIA | Hispanic |
| LH25 | Tumour | F | 65 | IIIB | Hispanic |
| LH26 | Tumour | M | 47 | IIIB | Hispanic |
| LH27 | Tumour | F | 46 | IIIB | Hispanic |
| LH28 | Tumour | M | 55 | IIIA | Hispanic |
| LH29 | Tumour | F | 71 | IIIA | Hispanic |
| LH30 | Tumour | M | 65 | IIIA | Hispanic |

**Foot notes:** H: Hispanic, NHW: Non-Hispanic White, LH: Local Hispanic, and NAT: Normal Adjacent Tissues.

**Supplementary Table 2:** Primer’s list

| Primers | Forward | Reverse |
| --- | --- | --- |
| BCL2L1 | AAGTCAACCACCAGCTCCC | CACTAACCAGAGACGAGACTCA |
| BCL2L12 | TCACCCGTGGACTTGAACTT | TGAGGTGTCATCTTGTAGCAGC |
| CCNB1 | CCAGAACCTGAGCCAGAACCT | GCTTGGAGAGGCAGTATCAACC |
| CDC25 | ATCTTCCTCCAGCATCTGAGTG | GGTGGAAAAATTCAAGGACAACACA |
| CDK1 | CATACCCATTGACTAACTATGGAAG | CCTGTAGTTTTGTGTCTACCCTT |
| CDK4 | GGGTGTAAGTGCCATCTGGT | CGAGATCTGAAGCCAGAGAACA |
| CHEK1 | GGTATTGGAATAACTCACAGGGA | CGAAATACTGTTGCCAAGCC |
| CSE1L | CGATGCAAATCTGCAAACACT | CAGTGGATAATTCTGATTTCCTTCA |
| ESPL1 | GGCTCCTCCTGCCACATC | GATCACAGGTCAGGGAAAGGA |
| FOXM1 | GGTCCAATGTCAAGTAGCGGT | CCAATGGCAAGGTCTCCTTCT |
| GAPDH | GACAACGAATTTGGCTACAGCA | GGTCTCTCTCTTCCTCTTGTGC |
| GLA | GAATCTCTTTGGGGAGCCATCC | GACTGCCAGGAAGAGCCAGAT |
| GPX1 | GGGAAACTCGCCTTGGTCTG | ACCTACGAGGGAGGAACACC |
| GPX2 | CAGGACGGACATACTTGAGACT | GAGAATGTGGCTTCGCTCTG |
| MCM10 | GGAGAGAACAACTTGCCTATCTG | GTAGCGCTCCTGCATCTCAG |
| MPV17 | ACTGGTATGCCCACTTTGAGG | GGAAGGCACATCGGCTCTAA |
| NQO1 | ATCCTGCCTGGAAGTTTAGGTC | CGCCTGGAGAATATTTGGGA |
| NUDT1 | AGTGCAAGAAGGAGAGACCATC | CACGAACTCAAACACGATCTGG |
| PDCD2L | CCATGGATCAGTTGCTTTCCC | GGGCAGGTCAAAAAGAGTGG |
| PMAIP1 | CTGCATTGTAATTGAGAGGAATGTG | TTCCATCTTCCGTTTCCAAGGG |
| PRDX1 | CTTCTGCCCTATCACTGAAAGCA | TCAGCCTGTCTGACTACAAAGG |
| PRDX4 | CTAGCCGCGACAACTCCG | GGCAGCAGGAACAGCAGTA |
| RRM2B | TACATCTCTGAGTGAACATTCTCG | GCAGCCAGTGATGGAATTGTA |
| SH3GLB1 | GCCCAGATGACTTACTATGCAC | CTGATGGTACAGGTGTCACAGA |
| SLC7A11 | TCTTCTGGTACAACTTCCAGTATT | GAGTCCCTGCGTATTATCTCTT |
| SOD2 | GTGACGTTCAGGTTGTTCACG | GACCTGCCCTACGACTACGG |
| TNFRSF12A | CTTCCAAGGTGTCTGGTTGC | CAGTTCCTTAGTCAGTGTCAGC |
| TRAF5 | GCGAGGAGAGTTTGACTCACT | GTCAGGTTTGAAGGTCTCCAT |
| TRIB3 | CTGTGTCGCTTTGTCTTCGC | GCTTGTCCCACAGGGAATCA |
| PPFIA1 | ATGATGTGCGAGGTGATGCC | GGCGGTCCCTTTCTTCTAGC |

**Supplementary Table 3:** Differentially expressed genes from both SW480 and Caco2 cell lines from transcriptomic analysis

| **Caco-2 DEGs** | | |
| --- | --- | --- |
| DEGs | Log2 fold change | p-adjusted value |
| ANKRD1 | 0.993833 | 0.012286 |
| AQP3 | -1.41969 | 1.03E-10 |
| ATF3 | 1.395377 | 1.13E-06 |
| CLCF1 | 1.029297 | 0.019358 |
| DNAJB9 | 1.119929 | 8.61E-09 |
| DPYSL3 | -1.11592 | 6.2E-07 |
| DUSP8 | 1.36561 | 0.000105 |
| FGF19 | 0.998549 | 0.017486 |
| GPR89B | 1.451921 | 3.21E-05 |
| IFIT1 | -2.15345 | 0.000156 |
| LIFR | 1.108374 | 0.006794 |
| MATN2 | -0.99515 | 0.004936 |
| MT2A | 2.698113 | 3.12E-17 |
| NGFR | 2.572154 | 3.22E-07 |
| NPPB | 1.790288 | 0.002401 |
| NPY1R | -1.57885 | 0.034781 |
| PTGS2 | 1.433735 | 3.56E-12 |
| SNHG12 | 1.223396 | 0.007099 |
| TMEM183B | -1.0955 | 0.006799 |
| TXNIP | 0.977554 | 0.004892 |
| ZNF286A | -0.96728 | 0.006778 |
| **SW480 DEGs** | | |
| DEGs | Log2 Fold Change | p-adjusted value |
| AK5 | -1.64937 | 5.81E-05 |
| CBX1 | -1.00335 | 2.7E-11 |
| CD36 | -1.0989 | 0.024536 |
| CLIC5 | -1.04831 | 0.031296 |
| DKK1 | -1.19004 | 8.72E-07 |
| DUSP15 | 1.176423 | 0.000453 |
| ESM1 | -1.79167 | 7.36E-05 |
| FAM122C | -1.05926 | 0.040637 |
| GLUL | -0.98673 | 4.14E-14 |
| HIST1H1C | -1.55891 | 1.64E-19 |
| HIST1H2AC | -1.28793 | 3.69E-08 |
| HIST1H2BC | -1.49159 | 0.02505 |
| HIST1H2BD | -1.04309 | 6.37E-05 |
| HIST1H2BJ | -1.80995 | 7.34E-10 |
| HIST1H2BK | -1.38422 | 2.4E-17 |
| HIST1H4I | -1.07669 | 4.48E-05 |
| HMOX1 | 1.796813 | 7.65E-15 |
| KLF2 | 1.289075 | 0.002098 |
| NT5E | -1.24257 | 0.001336 |
| PARM1 | -1.07264 | 0.013439 |
| PCTP | -1.0683 | 3.68E-05 |
| RASGRP3 | -1.16961 | 0.00051 |
| SEC61A1 | -0.96439 | 9.1E-18 |
| SEMA3C | -1.0549 | 0.000412 |
| SLC30A2 | 1.057649 | 0.040337 |
| SLC9A3 | 1.379976 | 6.33E-05 |
| TMEM45A | -0.98734 | 0.00678 |
| TNFSF9 | 1.055634 | 0.03741 |
| U2AF1 | -2.3836 | 4.64E-11 |
| UXS1 | -1.0886 | 1.98E-11 |
| ZNF26 | -1.00879 | 0.009046 |
| ZNF30 | -1.2373 | 0.000377 |
| ZNF302 | -1.07174 | 4.3E-08 |
| ZNF469 | 1.32969 | 4.48E-05 |
| ZNF594 | -1.14948 | 0.000442 |
| ZNF780A | -1.43379 | 1.44E-07 |
| ZNF780B | -1.1596 | 3.12E-05 |
| ZNF792 | -0.97342 | 0.015124 |

**Supplementary Table 4:** Differentially expressed genes from Caco2 cell lines from proteomic analysis

| **Protein Name** | **Gene Symbol** | **UNIPROT ID** | **log2Foldchange** | **pvalue** | **Molecular Weight** |
| --- | --- | --- | --- | --- | --- |
| Protein AAR2 homolog OS=Homo sapiens OX=9606 GN=AAR2 PE=1 SV=1 | AAR2 | A0A7P0TA70 | 1.584962501 | 0.070484 | 50 kDa |
| Complex I assembly factor ACAD9, mitochondrial OS=Homo sapiens OX=9606 GN=ACAD9 PE=1 SV=1 | ACAD9 | Q9H845 | 2.169925001 | 0.072827 | 69 kDa |
| Acireductone dioxygenase OS=Homo sapiens OX=9606 GN=ADI1 PE=1 SV=1 | ADI1 | Q9BV57 | -2.807354922 | 0.07645 | 21 kDa |
| Protein argonaute-2 OS=Homo sapiens OX=9606 GN=AGO2 PE=1 SV=3 | AGO2 | Q9UKV8 | 3 | 0.007763 | 97 kDa |
| Aldo-keto reductase family 1 member C2 OS=Homo sapiens OX=9606 GN=AKR1C2 PE=1 SV=3 | AKR1C2 | P52895 | 1 | 0.007763 | 37 kDa |
| Probable bifunctional dTTP/UTP pyrophosphatase/methyltransferase protein OS=Homo sapiens OX=9606 GN=ASMTL PE=1 SV=3 | ASMTL | O95671 | 1.584962501 | 0.07418 | 69 kDa |
| Cluster of V-type proton ATPase subunit OS=Homo sapiens OX=9606 GN=ATP6V0D1 PE=1 SV=1+1 | ATP6V0D1 | F5GYQ1 (+1) | 1.584962501 | 0.070484 |  |
| V-type proton ATPase subunit C 1 OS=Homo sapiens OX=9606 GN=ATP6V1C1 PE=1 SV=4 | ATP6V1C1 | P21283 | -1.807354922 | 0.049372 | 44 kDa |
| Aurora kinase A OS=Homo sapiens OX=9606 GN=AURKA PE=1 SV=3 | AURKA | O14965 | 2.584962501 | 0.03775 |  |
| Cluster of Aurora kinase A OS=Homo sapiens OX=9606 GN=AURKA PE=1 SV=3+2 | AURKB (+1) | A3KFJ0 (+2) | 1.807354922 | 0.049372 |  |
| Brain-specific angiogenesis inhibitor 1-associated protein 2 OS=Homo sapiens OX=9606 GN=BAIAP2 PE=1 SV=1 | BAIAP2 | I3L4C2 | -1.847996907 | 0.054735 | 61 kDa |
| Probable 18S rRNA (guanine-N(7))-methyltransferase OS=Homo sapiens OX=9606 GN=BUD23 PE=1 SV=2 | BUD23 | O43709 | 2.169925001 | 0.05844 | 32 kDa |
| Splicing factor C9orf78 OS=Homo sapiens OX=9606 GN=C9orf78 PE=1 SV=1 | C9orf78 | Q9NZ63 | -1 | 0.053455 | 34 kDa |
| Cyclin-K OS=Homo sapiens OX=9606 GN=CCNK PE=1 SV=2 | CCNK | O75909 | 1 | 0.035099 | 64 kDa |
| CDK5 regulatory subunit-associated protein 3 OS=Homo sapiens OX=9606 GN=CDK5RAP3 PE=1 SV=2 | CDK5RAP3 | Q96JB5 | -1.192645078 | 0.057638 | 57 kDa |
| CDKN2AIP N-terminal-like protein OS=Homo sapiens OX=9606 GN=CDKN2AIPNL PE=1 SV=1 | CDKN2AIPNL | Q96HQ2 | -1.736965594 | 0.019804 | 13 kDa |
| Charged multivesicular body protein 1a OS=Homo sapiens OX=9606 GN=CHMP1A PE=1 SV=1 | CHMP1A | F8VUA2 | 2 | 0.07645 | 20 kDa |
| Group of Chromatin target of PRMT1 OS=Homo sapiens OX=9606 GN=CHTOP PE=4 SV=1+1 | CHTOP | A0AA34QVV3 (+1) | 2.321928095 | 0.047421 |  |
| Clathrin light chain B OS=Homo sapiens OX=9606 GN=CLTB PE=1 SV=1 | CLTB | P09497 | -1.415037499 | 0.082442 | 25 kDa |
| CCR4-NOT transcription complex subunit 11 OS=Homo sapiens OX=9606 GN=CNOT11 PE=1 SV=1 | CNOT11 | Q9UKZ1 | -1.807354922 | 0.02411 | 55 kDa |
| Cullin 4A OS=Homo sapiens OX=9606 GN=CUL4A PE=1 SV=1 | CUL4A | A0A0A0MR50 | 1.087462841 | 0.062147 | 78 kDa |
| DNA damage inducible 1 homolog 2 OS=Homo sapiens OX=9606 GN=DDI2 PE=4 SV=1 | DDI2 | A0AA34QVV2 | 1.280107919 | 0.049372 | 47 kDa |
| Dihydropyrimidinase-related protein 3 OS=Homo sapiens OX=9606 GN=DPYSL3 PE=1 SV=1 | DPYSL3 | Q14195 | -1.321928095 | 0.013236 | 62 kDa |
| Developmentally-regulated GTP-binding protein 2 OS=Homo sapiens OX=9606 GN=DRG2 PE=1 SV=1 | DRG2 | P55039 | 1.807354922 | 0.02411 | 41 kDa |
| Group of Probable RNA-binding protein EIF1AD (Fragment) OS=Homo sapiens OX=9606 GN=EIF1AD PE=1 SV=1+2 | EIF1AD | E9PLI6 (+2) | -1.321928095 | 0.013236 |  |
| Ephrin type-A receptor 2 OS=Homo sapiens OX=9606 GN=EPHA2 PE=1 SV=2 | EPHA2 | P29317 | 3.169925001 | 0.024817 | 108 kDa |
| Epsin-1 OS=Homo sapiens OX=9606 GN=EPN1 PE=1 SV=2 | EPN1 | Q9Y6I3 | -1.415037499 | 0.067197 | 60 kDa |
| Ferritin OS=Homo sapiens OX=9606 GN=FTH1 PE=1 SV=1 | FTH1 | G3V192 | 2.321928095 | 0.047421 | 18 kDa |
| RNA-binding protein FXR1 OS=Homo sapiens OX=9606 GN=FXR1 PE=1 SV=1 | FXR1 | A0A8V8TMQ1 | 1.169925001 | 0.082442 | 73 kDa |
| Glutamine synthetase OS=Homo sapiens OX=9606 GN=GLUL PE=1 SV=1 | GLUL | A0A2R8YDT1 | -1.906890596 | 0.05226 | 57 kDa |
| Guanine nucleotide-binding protein G(I)/G(S)/G(O) subunit gamma-10 OS=Homo sapiens OX=9606 GN=GNG10 PE=1 SV=1 | GNG10 | P50151 | -2 | 0.07645 | 7 kDa |
| Golgi membrane protein 1 OS=Homo sapiens OX=9606 GN=GOLM1 PE=1 SV=1 | GOLM1 | Q8NBJ4 | -1.584962501 | 0.057191 | 45 kDa |
| Cluster of HCLS1-associated protein X-1 OS=Homo sapiens OX=9606 GN=HAX1 PE=1 SV=1+1 | HAX1 | A0A8V8TLX9 (+1) | -2.584962501 | 0.03775 |  |
| Hexokinase-2 OS=Homo sapiens OX=9606 GN=HK2 PE=1 SV=2 | HK2 | P52789 | 2.321928095 | 0.047421 | 102 kDa |
| Hook microtubule tethering protein 3 OS=Homo sapiens OX=9606 GN=HOOK3 PE=1 SV=2 | HOOK3 | H0YDM4 | -1.584962501 | 0.07418 | 83 kDa |
| Cation-independent mannose-6-phosphate receptor OS=Homo sapiens OX=9606 GN=IGF2R PE=1 SV=3 | IGF2R | P11717 | 1.473931188 | 0.01663 | 274 kDa |
| Kinesin-like protein OS=Homo sapiens OX=9606 GN=KIF11 PE=1 SV=1 | KIF11 | A0A7I2V3A9 | 1 | 0.047421 | 112 kDa |
| Lysine-rich nucleolar protein 1 OS=Homo sapiens OX=9606 GN=KNOP1 PE=1 SV=1 | KNOP1 | Q1ED39 | 1.807354922 | 0.02411 | 52 kDa |
| Galectin-2 OS=Homo sapiens OX=9606 GN=LGALS2 PE=1 SV=3 | LGALS2 | P05162 | -1.280107919 | 0.095061 | 15 kDa |
| Lysophosphatidylcholine acyltransferase 2 OS=Homo sapiens OX=9606 GN=LPCAT2 PE=1 SV=1 | LPCAT2 | Q7L5N7 | -2.584962501 | 0.03775 | 60 kDa |
| Leucine-rich repeat flightless-interacting protein 2 OS=Homo sapiens OX=9606 GN=LRRFIP2 PE=1 SV=1 | LRRFIP2 | Q9Y608 | -1.807354922 | 0.02411 | 82 kDa |
| Complex III assembly factor LYRM7 OS=Homo sapiens OX=9606 GN=LYRM7 PE=1 SV=1 | LYRM7 | Q5U5X0 | -1.584962501 | 0.057191 | 12 kDa |
| Large ribosomal subunit protein bL28m OS=Homo sapiens OX=9606 GN=MRPL28 PE=1 SV=4 | MRPL28 | Q13084 | 1.169925001 | 0.082442 | 30 kDa |
| Large ribosomal subunit protein mL37 OS=Homo sapiens OX=9606 GN=MRPL37 PE=1 SV=2 | MRPL37 | Q9BZE1 | 2.807354922 | 0.013236 | 48 kDa |
| Mitochondrial ribosomal protein L48 OS=Homo sapiens OX=9606 GN=MRPL48 PE=1 SV=1 | MRPL48 | F5H702 | 1.584962501 | 0.07418 | 13 kDa |
| NADH dehydrogenase [ubiquinone] 1 alpha subcomplex assembly factor 2 OS=Homo sapiens OX=9606 GN=NDUFAF2 PE=1 SV=1 | NDUFAF2 | Q8N183 | -1 | 0.053455 | 20 kDa |
| NADH dehydrogenase [ubiquinone] 1 alpha subcomplex assembly factor 3 OS=Homo sapiens OX=9606 GN=NDUFAF3 PE=1 SV=1 | NDUFAF3 | Q9BU61 | 1.415037499 | 0.082442 | 20 kDa |
| Nucleolar complex protein 2 homolog OS=Homo sapiens OX=9606 GN=NOC2L PE=1 SV=4 | NOC2L | Q9Y3T9 | 1.874469118 | 0.015433 | 85 kDa |
| Vesicle-fusing ATPase OS=Homo sapiens OX=9606 GN=NSF PE=1 SV=1 | NSF | I3L0N3 | 1 | 0.047421 | 82 kDa |
| Nucleobindin-2 OS=Homo sapiens OX=9606 GN=NUCB2 PE=1 SV=3 | NUCB2 | P80303 | -1.201633861 | 0.0107 | 50 kDa |
| ADP-ribose pyrophosphatase, mitochondrial OS=Homo sapiens OX=9606 GN=NUDT9 PE=1 SV=1 | NUDT9 | Q9BW91 | -2.169925001 | 0.035653 | 39 kDa |
| tRNA N6-adenosine threonylcarbamoyltransferase OS=Homo sapiens OX=9606 GN=OSGEP PE=1 SV=1 | OSGEP | Q9NPF4 | 2.807354922 | 0.013236 | 36 kDa |
| PDZ and LIM domain protein 7 OS=Homo sapiens OX=9606 GN=PDLIM7 PE=1 SV=1 | PDLIM7 | Q9NR12 | 1.415037499 | 0.082442 | 50 kDa |
| PDZ domain-containing protein 11 OS=Homo sapiens OX=9606 GN=PDZD11 PE=1 SV=2 | PDZD11 | Q5EBL8 | -1.584962501 | 0.057191 | 16 kDa |
| Mitochondrial-processing peptidase subunit alpha OS=Homo sapiens OX=9606 GN=PMPCA PE=1 SV=2 | PMPCA | Q10713 | 1 | 0.048646 | 58 kDa |
| Cluster of DNA polymerase OS=Homo sapiens OX=9606 GN=POLD1 PE=1 SV=1+1 | POLD1 | M0R2B7 (+1) | 1.192645078 | 0.028596 |  |
| Ribonucleases P/MRP protein subunit POP1 OS=Homo sapiens OX=9606 GN=POP1 PE=1 SV=2 | POP1 | Q99575 | 1.584962501 | 0.067197 | 115 kDa |
| Cluster of POU domain protein (Fragment) OS=Homo sapiens OX=9606 GN=POU2F1 PE=1 SV=1+1 | POU2F1 | A0A0C4DG88 (+1) | -2.169925001 | 0.035653 |  |
| Amidophosphoribosyltransferase OS=Homo sapiens OX=9606 GN=PPAT PE=1 SV=1 | PPAT | Q06203 | 2.321928095 | 0.047421 | 57 kDa |
| PTPRF interacting protein alpha 1 (Fragment) OS=Homo sapiens OX=9606 GN=PPFIA1 PE=1 SV=1 | PPFIA1 | A0A2R8Y7R9 | -1.263034406 | 0.035653 | 139 kDa |
| Protein phosphatase 1 regulatory subunit 12A OS=Homo sapiens OX=9606 GN=PPP1R12A PE=1 SV=1 | PPP1R12A | O14974 | -1.584962501 | 0.035653 | 115 kDa |
| Small ribosomal subunit protein mS39 OS=Homo sapiens OX=9606 GN=PTCD3 PE=1 SV=3 | PTCD3 | Q96EY7 | 1.137503524 | 0.013236 | 79 kDa |
| Pseudouridylate synthase 7 homolog OS=Homo sapiens OX=9606 GN=PUS7 PE=1 SV=2 | PUS7 | Q96PZ0 | 1 | 0.038561 | 75 kDa |
| Paxillin OS=Homo sapiens OX=9606 GN=PXN PE=1 SV=1 | PXN | A0A1B0GTU4 | -1 | 0.039872 | 116 kDa |
| Queuine tRNA-ribosyltransferase accessory subunit 2 OS=Homo sapiens OX=9606 GN=QTRT2 PE=1 SV=1 | QTRT2 | Q9H974 | 2.807354922 | 0.013236 | 47 kDa |
| Group of Rab3 GTPase-activating protein catalytic subunit OS=Homo sapiens OX=9606 GN=RAB3GAP1 PE=1 SV=1+1 | RAB3GAP1 | A0A8J9AUI2 (+1) | -1.378511623 | 0.047421 |  |
| Rap1 GTPase-GDP dissociation stimulator 1 OS=Homo sapiens OX=9606 GN=RAP1GDS1 PE=1 SV=3 | RAP1GDS1 | P52306 | 2.321928095 | 0.047421 | 66 kDa |
| Regulator of G-protein signaling 10 OS=Homo sapiens OX=9606 GN=RGS10 PE=1 SV=3 | RGS10 | O43665 | -2.584962501 | 0.082442 | 21 kDa |
| E3 ubiquitin-protein ligase RING1 OS=Homo sapiens OX=9606 GN=RING1 PE=1 SV=2 | RING1 | Q06587 | 2.584962501 | 0.03775 | 42 kDa |
| Ribosomal protein S6 kinase beta-2 OS=Homo sapiens OX=9606 GN=RPS6KB2 PE=1 SV=2 | RPS6KB2 | Q9UBS0 | 1.415037499 | 0.082442 | 53 kDa |
| Splicing factor 3B subunit 6 OS=Homo sapiens OX=9606 GN=SF3B6 PE=1 SV=1 | SF3B6 | Q9Y3B4 | 1 | 0.078155 | 15 kDa |
| STE20-like serine/threonine-protein kinase OS=Homo sapiens OX=9606 GN=SLK PE=1 SV=1 | SLK | Q9H2G2 | -2.321928095 | 0.004813 | 143 kDa |
| Kunitz-type protease inhibitor 2 OS=Homo sapiens OX=9606 GN=SPINT2 PE=1 SV=2 | SPINT2 | O43291 | -1 | 0.004813 | 28 kDa |
| Small ubiquitin-related modifier 1 OS=Homo sapiens OX=9606 GN=SUMO1 PE=1 SV=1 | SUMO1 | P63165 | -1.906890596 | 0.053271 | 12 kDa |
| Ubiquitin-conjugating enzyme E2 E2 OS=Homo sapiens OX=9606 GN=UBE2E2 PE=1 SV=1 | UBE2E2 | Q96LR5 | 2.807354922 | 0.013236 | 22 kDa |
| Small subunit processome component 20 homolog OS=Homo sapiens OX=9606 GN=UTP20 PE=1 SV=3 | UTP20 | O75691 | 1 | 0.078155 | 318 kDa |
| Cytoplasmic 60S subunit biogenesis factor ZNF622 OS=Homo sapiens OX=9606 GN=ZNF622 PE=1 SV=1 | ZNF622 | Q969S3 | 1 | 0.047421 | 54 kDa |
| Zinc finger protein 638 OS=Homo sapiens OX=9606 GN=ZNF638 PE=1 SV=2 | ZNF638 | Q14966 | 1.222392421 | 0.057191 | 221 kDa |
